# Transcriptome dissection of the Zeb2-associated network reveals a role for *NRPS15* in zearalenone biosynthesis in *Fusarium graminearum*

**DOI:** 10.64898/2026.10.23.761285

**Authors:** Nahyun Lee, Jae Woo Han, Jung-Eun Kim, Jiyoung Shin, Ae Ran Park, Yin-Won Lee, Hun Kim, Jiyeun Park, Hokyoung Son

## Abstract

Fungal secondary metabolites, including mycotoxins, play important roles in fungal biology and represent major concerns for food and feed safety. Zearalenone (ZEA) is an estrogenic polyketide mycotoxin produced by several *Fusarium* species, yet the genetic network governing its production remains poorly characterized. Here, we investigated genes associated with ZEA production in *Fusarium graminearum* using two complementary transcriptomic approaches. Time-resolved transcriptome analysis revealed transcriptional changes preceding detectable ZEA accumulation. Temporal clustering further revealed coordinated changes in primary metabolism, transcriptional regulation, carbohydrate utilization, and stress- and transport-associated pathways during ZEA-inducing cultivation. Genotype-based transcriptome analysis of *ZEB2* deletion and overexpression strains subsequently identified Zeb2-responsive genes. Functional screening identified *NRPS15*, encoding a previously uncharacterized nonribosomal peptide synthetase-like protein adjacent to the ZEA biosynthetic cluster, as an important contributor to ZEA production. Deletion of *NRPS15* markedly reduced ZEA production and expression of *PKS4*, *PKS13*, *ZEB1*, and *ZEB2*. *NRPS15* expression did not restore ZEA production in the absence of Zeb2. *NRPS15* orthologs were conserved in several ZEA-producing *Fusarium* species, and deletion of *NRPS15* substantially reduced ZEA production in *Fusarium culmorum*. These findings expand the functional genetic context of ZEA production beyond the canonical four-gene biosynthetic cluster.

## 1. Introduction

Fungal secondary metabolites play important roles in fungal ecology, development, pathogenicity, and interactions with other organisms, while many also have substantial impacts on agriculture, food safety, and human and animal health (Keller, 2019; Macheleidt et al., 2016). Fungal secondary metabolism is commonly organized within biosynthetic gene clusters, in which genes required for the production of a specific metabolite are physically linked in the genome. Such clustering can facilitate coordinated gene expression and regulation and may provide evolutionary advantages by maintaining functionally related genes as a unit (Rokas et al., 2018; Wisecaver and Rokas, 2015). Many biosynthetic gene clusters also encode pathway-specific transcription factors that coordinate the expression of their associated biosynthetic genes. Importantly, the influence of these regulators may extend beyond their associated biosynthetic gene clusters (Kwon et al., 2021; Shostak et al., 2020). For example, AflR-responsive genes have been identified outside the aflatoxin biosynthetic cluster in *Aspergillus parasiticus*, whereas Tri6 in *Fusarium graminearum* influences genes involved in primary metabolism, transport, signaling, and additional secondary metabolic pathways beyond the trichothecene cluster (Nasmith et al., 2011; Shostak et al., 2020). Thus, pathway-specific regulators can participate in broader transcriptional networks that connect secondary metabolite production with diverse cellular processes.

Recent studies have also shown that genetic factors influencing fungal secondary metabolism are not limited to well-characterized pathway-specific regulators or the initially defined core biosynthetic genes. Biosynthetic loci frequently contain neighboring genes encoding tailoring enzymes, transporters, regulatory proteins, and other accessory factors, and functional analyses have revealed that such genes can make unexpected contributions to metabolite production (Chang et al., 2009; Sieber et al., 2014). In *Fusarium fujikuroi*, genes neighboring the core polyketide synthase gene *BIK1* were shown to encode enzymes, regulators, and a transporter required for efficient bikaverin production (Wiemann et al., 2009). Similarly, analysis of the asperfuranone pathway in *Aspergillus nidulans* demonstrated that several genes adjacent to the core polyketide synthases contribute substantially to metabolite biosynthesis (Chiang et al., 2009). More broadly, comparative genomic studies have shown that the organization and synteny of secondary-metabolite loci can vary among related fungal species through gene gain, loss, or rearrangement (Chang et al., 2009; Sieber et al., 2014). Biosynthetic loci do not always conform to simple, contiguous genomic boundaries. In *Aspergillus fumigatus*, genes involved in the biosynthesis of two chemically distinct secondary metabolites, fumagillin and pseurotin, are interspersed within the same genomic region rather than organized as two clearly separated biosynthetic clusters (Wiemann et al., 2013). Collectively, these findings demonstrate that neighboring genes can form an extended functional unit with core biosynthetic genes and that the functional boundaries of fungal secondary-metabolite loci may be broader and more complex than initially predicted.

*F. graminearum* is a major fungal pathogen of cereal crops and produces multiple mycotoxins that can accumulate in infected cereals. Among these, zearalenone (ZEA) is a polyketide-derived, nonsteroidal estrogenic mycotoxin produced by several *Fusarium* species, including *F. graminearum*, *F. pseudograminearum*, and *F. culmorum*. ZEA frequently contaminates cereals and cereal-derived products and can cause estrogenic and reproductive disorders, particularly in livestock (Katzenellenbogen et al., 1979; Zhen et al., 2024). Understanding the molecular mechanisms governing ZEA production is therefore important for understanding secondary metabolism in *Fusarium* species and for developing strategies to reduce mycotoxin contamination in food and feed.

The core biosynthetic pathway for ZEA has been characterized in *F. graminearum*. The ZEA biosynthetic gene cluster contains two polyketide synthase genes, *PKS4* and *PKS13*, together with *ZEB1* and *ZEB2*. Pks4 and Pks13 cooperatively generate the polyketide backbone, whereas Zeb1 catalyzes an oxidation step associated with conversion of β-zearalenol to ZEA (Gaffoor and Trail, 2006; Kim et al., 2018; Kim et al., 2005). Zeb2 functions as a pathway-specific transcription factor that controls the expression of the ZEA biosynthetic genes. Previous studies further revealed an unusual autoregulatory mechanism governing *ZEB2* expression involving two isoforms, Zeb2L and Zeb2S, generated through alternative promoter usage (Park et al., 2015). Zeb2L contains an N-terminal basic leucine zipper DNA-binding domain, whereas Zeb2S lacks this region and modulates Zeb2L activity through heterodimer formation. ZEA itself can induce expression of both isoforms, establishing a feedback mechanism that contributes to regulation of pathway activity. These studies establish Zeb2 as a central regulator of ZEA biosynthesis; however, whether its regulatory influence extends beyond the canonical ZEA biosynthetic cluster remains poorly understood.

In this study, we sought to define the broader Zeb2-associated network involved in ZEA production in *F. graminearum*. We combined time-resolved transcriptome profiling with comparative RNA-seq analyses of the wild-type strain and *ZEB2* mutants to identify genes exhibiting *ZEB2*-dependent expression patterns. Functional screening of these candidates identified *NRPS15*, which encodes a previously uncharacterized nonribosomal peptide synthetase (NRPS)-like protein located adjacent to the ZEA biosynthetic gene cluster, as an important contributor to ZEA production. These findings reveal an unexpected link between an NRPS-like protein and regulation of the ZEA pathway and suggest that the functional genetic context underlying ZEA production extends beyond the canonical biosynthetic gene cluster.

## 2. Materials and Methods

### 2.1. Fungal strains and culture conditions

The wild-type strain used in this study was *F. graminearum* Z-3639 (Bowden and Leslie, 1999), and all strains used in this study are listed in Table S1. The *F. culmorum* strain used in this study was obtained from the Korean Agricultural Culture Collection (accession no. KACC 48423). Culture media were prepared and fungal strains were cultured according to the *Fusarium* laboratory manual (Leslie and Summerell, 2006). For genomic DNA extraction, fungal strains were cultured in liquid complete medium (CM) at 25 °C for 5 days, after which the mycelia were harvested and lyophilized. Conidia were produced on yeast malt agar (YMA) or in carboxymethyl cellulose (CMC) medium (Harris, 2005), whereas starch–glutamate (SG) liquid medium was used for ZEA production and quantification (Park et al., 2015). All wild-type and mutant strains were cultured on CM at 25 °C and stored at −80 °C in 20% glycerol.

### 2.2. Phenotypic characterization and pathogenicity assay

To assess vegetative growth, each strain was cultured on CM and minimal medium (MM) for 4 days. For conidiation assays, fresh mycelial plugs grown on CM agar were inoculated into 20 mL of CMC medium and incubated at 25 °C with shaking at 150 rpm for 5 days. Conidia were harvested, and their concentrations were determined using a hemocytometer. To assess sexual development, strains were cultured on carrot agar for 5 days, after which aerial mycelia were removed by gently scraping the surface with 0.4 mL of sterile 2.5% Tween 60. The cultures were subsequently incubated under near-UV light for 8–9 days, and perithecium formation was assessed. Pathogenicity assays were performed using conidia produced in CMC medium. Conidia were harvested and adjusted to 1 × 10^5^ conidia/mL, and 10 µL of the conidial suspension was inoculated into spikelets of the wheat cultivar Eunpamil. Inoculated plants were maintained at >90% relative humidity for 3 days and subsequently transferred to greenhouse conditions. Disease development was evaluated 3 weeks after inoculation by counting the number of symptomatic spikelets.

### 2.3. ZEA-inducing culture conditions and quantification

ZEA production was induced according to a previously described protocol (Kim et al., 2018). Fresh mycelia obtained after 2 days of germination were transferred to 20 mL SG medium and incubated at 24 °C for the indicated periods depending on the experiment. Cultures were then filtered through Miracloth, and the resulting culture filtrates were used for ZEA quantification, whereas the harvested mycelia were subjected to RNA extraction and transcriptome analysis. For ZEA extraction, 1 mL of culture filtrate was mixed with 4 mL of ethyl acetate and incubated for 30 min at 25 °C. The organic phase was concentrated using a SpeedVac (Thermo Fisher Scientific, Waltham, MA, USA) and subsequently reconstituted in methanol–water (70:30, v/v). ZEA was quantified using a Prominence HPLC system (Shimadzu, Japan) equipped with a reverse-phase C18 column. The mobile phase consisted of 70% aqueous methanol and was delivered at a flow rate of 1 mL/min.

### 2.4. RNA extraction and quantitative reverse transcription PCR

Total RNA was extracted from harvested fungal mycelia using the Easy-Spin Total RNA Extraction Kit (iNtRON Biotechnology, Seoul, Republic of Korea). RNA concentration and purity were assessed with a NanoDrop spectrophotometer (Thermo Fisher Scientific, Waltham, MA, USA). First-strand cDNA was synthesized using the SuperScript III First-Strand Synthesis System (Invitrogen, Carlsbad, CA, USA). Quantitative reverse transcription PCR was conducted on a CFX Real-Time PCR System (Bio-Rad, Hercules, CA, USA) with iTaq Universal SYBR Green Supermix (Bio-Rad). Three or four independent biological replicates were included for each strain and experimental condition. Relative transcript levels were normalized against the endogenous cyclophilin gene (*CYP1*). The primers used for quantitative reverse transcription PCR (RT-qPCR) are listed in Table S2.

### 2.5. RNA-seq data processing and differential expression analysis

Raw sequence reads were subjected to quality control using FastQC (v0.11.9) and low-quality bases and adapter sequences were trimmed using fastp (v1.3.3). The resulting data were aligned to the *F. graminearum* PH-1 reference genome using the STAR aligner with default parameters. Differentially expressed genes (DEGs) were identified using the DESeq2 R package (v1.50.2). Statistical significance was determined based on an adjusted p-value (FDR) < 0.05 and a minimum absolute log_2_ fold-change of 1. Functional enrichment analysis, including Gene Ontology (GO) and Kyoto Encyclopedia of Genes and Genomes (KEGG), was conducted using clusterProfiler R package (v4.18.4). Terms and pathways with an adjusted p-value < 0.05 were considered significantly enriched.

### 2.6. Soft clustering analysis (Mfuzz)

Soft clustering was performed using the Mfuzz (v2.70.0) R package. Raw counts were first normalized using the variance stabilizing transformation (VST) in DESeq2. For clustering, we used a union set of 7,569 DEGs identified across all time points. Expression values were averaged across biological replicates at each time point to generate a representative expression matrix. The fuzzifier parameter *m* was objectively estimated using the mestimate function, and the number of clusters was set to *k* = 6. Genes were assigned to specific clusters based on their membership values.

### 2.7. Genetic manipulation of fungal strains

Gene deletion constructs were generated using the double-joint PCR and split-marker recombination methods (Catlett et al., 2003; Yu et al., 2004). The 5′ and 3′ flanking regions of each target gene were amplified from wild type genomic DNA, whereas the geneticin resistance cassette (*GEN*) was amplified from plasmid pII99 (Namiki et al., 2001). The three fragments were fused by double-joint PCR, and the final replacement construct was amplified using nested primers. The resulting amplicons were introduced into wild type protoplasts by polyethylene glycol-mediated transformation according to a previously described protocol (Han et al., 2007). Correct replacement of the target gene was verified by Southern hybridization using a probe corresponding to one of the flanking regions (Fig. S1).

*NRPS15* complementation and overexpression constructs were assembled using a yeast gap-repair cloning approach (Zhou et al., 2011). For complementation, a fragment containing the native *NRPS15* promoter and open reading frame was amplified. For overexpression, the *NRPS15* open reading frame was assembled downstream of the constitutive RP27 promoter. The respective fragments were co-transformed with XhoI-linearized pDL2 into *Saccharomyces cerevisiae* strain PJ69-4A using an alkali-cation yeast transformation kit. Recombinant plasmids were recovered from the transformed yeast cells and subsequently introduced into *Escherichia coli* DH10B for propagation. After verification by PCR and sequencing, the confirmed complementation and overexpression vectors were introduced into Δ*nrps15* protoplasts by PEG-mediated transformation.

### 2.8. Heterologous expression of *NRPS15* in *S. cerevisiae*

The *NRPS15* open reading frame was amplified using primers containing HindIII and XhoI restriction sites at the 5′ and 3′ ends, respectively. The amplified fragment was digested with HindIII and XhoI and ligated into the corresponding sites of the pYES2 expression vector. The resulting recombinant plasmid was introduced into *S. cerevisiae* strain BY4741, whereas yeast transformed with the empty pYES2 vector was used as a control. Correct insertion of *NRPS15* was verified by PCR and sequencing.

Expression of *NRPS15* in the transformed yeast strain was confirmed by RT-qPCR. Yeast cultures were adjusted to an OD_600_ of 0.5, harvested by centrifugation, and ground to a fine powder in liquid nitrogen. Total RNA was extracted from the resulting cell powder using the Easy-Spin Total RNA Extraction Kit, and cDNA synthesis and RT-qPCR were performed as described above.

### 2.9. Metabolite extraction and profiling of *NRPS15*-expressing yeast

Each strain was precultured in YPD medium for 24 h, and the preculture was subsequently inoculated at 1% (v/v) into 100 mL of YP medium containing galactose as the carbon source in a 500-mL baffled Erlenmeyer flask. Cultures were incubated at 30 °C and 150 rpm for 3 days. Following cultivation, the cultures were centrifuged at 10,000 × g for 20 min to separate the culture supernatants from the cell pellets.

For intracellular metabolite extraction, each cell pellet was resuspended in 100 mL of methanol–acetone (9:1, v/v), sonicated for 1 h, and incubated overnight at room temperature. The resulting extracts and culture supernatants were filtered through 0.2-µm polytetrafluoroethylene syringe filters. Extracellular and intracellular metabolites were initially analyzed by HILIC–HPLC and UPLC–ESI–MS. HILIC–HPLC analysis was performed using a Shimadzu LC-40D system equipped with a YMC-Triart Diol-HILIC column (150 × 4.6 mm, 5 µm; YMC Co., Ltd., Kyoto, Japan) under isocratic conditions with acetonitrile–water (90:10, v/v), and chromatograms were monitored at 210 nm. UPLC–ESI–MS analysis was performed using a Waters ACQUITY UPLC H-Class PLUS system coupled to a Waters SQ Detector 2 (Waters Corporation, Milford, MA, USA) with an ACQUITY UPLC BEH C18 column (2.1 × 50 mm, 1.7 µm; Waters Corporation). Samples were analyzed in positive-ion mode using water and acetonitrile containing 0.1% formic acid as mobile phases. Full-scan spectra acquired throughout the chromatographic run were combined for comparison of composite ion profiles.

To broaden the range of extracellular metabolites recovered, culture supernatants from the *NRPS15*-expressing strain and empty-vector control were sequentially extracted twice with equal volumes of ethyl acetate (EtOAc) and *n*-butanol (*n*-BuOH). The solvent fractions were separately pooled, concentrated to dryness, and analyzed by reverse-phase HPLC and GC–MS. RP-HPLC analysis was performed using a Shimadzu Prominence LC-20AR system equipped with a Phenomenex Luna C18(2) column (250 × 4.6 mm, 5 µm; Phenomenex, Torrance, CA, USA), with a linear gradient of 20–100% methanol containing 0.1% formic acid over 20 min. Chromatograms were monitored at 230 nm.

For GC–MS analysis, dried EtOAc and *n*-BuOH extracts were subjected to methoximation and trimethylsilylation using methoxyamine hydrochloride and BSTFA containing 1% TMCS. Derivatized samples were analyzed using a Shimadzu GCMS-QP2020 system (Shimadzu, Kyoto, Japan) equipped with a Rxi-5ms capillary column (30 m × 0.25 mm internal diameter, 0.25 µm film thickness; Restek, Bellefonte, PA, USA). Mass spectra were acquired in electron-ionization mode at 70 eV over an *m*/*z* range of 50–700.

### 2.10. Comparative sequence, domain, and genomic context analysis of *NRPS15*

The amino acid sequence of *F. graminearum NRPS15* was used as a query for BLASTp searches to identify homologs in other *Fusarium* species and to assess sequence conservation. The *NRPS15* homologs analyzed included XP_009258254.1 from *F. pseudograminearum*, RGP69730.1 from *F. sporotrichioides*, and PTD10916 from *F. culmorum*. Conserved domains and NRPS domain organization were examined using the NCBI Conserved Domain Database (CDD). The predicted adenylation domains identified by CDD were further analyzed using PARAS (Terlouw et al., 2026) to predict substrate specificity. antiSMASH (Blin et al., 2023) was used to examine biosynthetic gene cluster organization, the genomic context of *NRPS15*, and its positional relationship with the ZEA biosynthetic gene cluster in the respective *Fusarium* genomes.

### 2.11. Statistical analysis

All experiments were performed with at least three independent biological replicates unless otherwise indicated. Data are presented as the mean ± standard deviation. Statistical analyses were performed using GraphPad Prism (GraphPad Software, Boston, MA, USA). Comparisons between two groups were performed using two-tailed Student’s t-tests. For comparisons among multiple strains, one-way analysis of variance (ANOVA) followed by an appropriate multiple-comparison test was performed. Differences were considered statistically significant at *p* < 0.05. Significance levels are indicated as follows: \**p* < 0.05, \*\**p* < 0.01, \*\*\**p* < 0.001, and \*\*\*\**p* < 0.0001; ns, not significant.

## 3. Results

### 3.1. Temporal transcriptomic profiling of *F. graminearum* under zearalenone-inducing conditions

To investigate the transcriptional dynamics of *F. graminearum* during zearalenone production, time-course RNA-seq analysis was performed using samples collected from SG medium at 6 h, 2 d, 4 d, 6 d, 8 d, and 10 d. Principal component analysis (PCA) was employed to assess global transcriptomic variation and revealed that the first two components (PC1 and PC2) accounted for 66% and 28% of the total variance, respectively (Fig. 1A). Samples collected at 6 h were clearly separated from those collected at later time points along PC1. A pronounced transcriptomic transition occurred between 6 h and 2 d, whereas samples collected from 6 d to 10 d clustered more closely, indicating relatively limited global transcriptional changes during the later stages of cultivation.

**Fig. 1.**
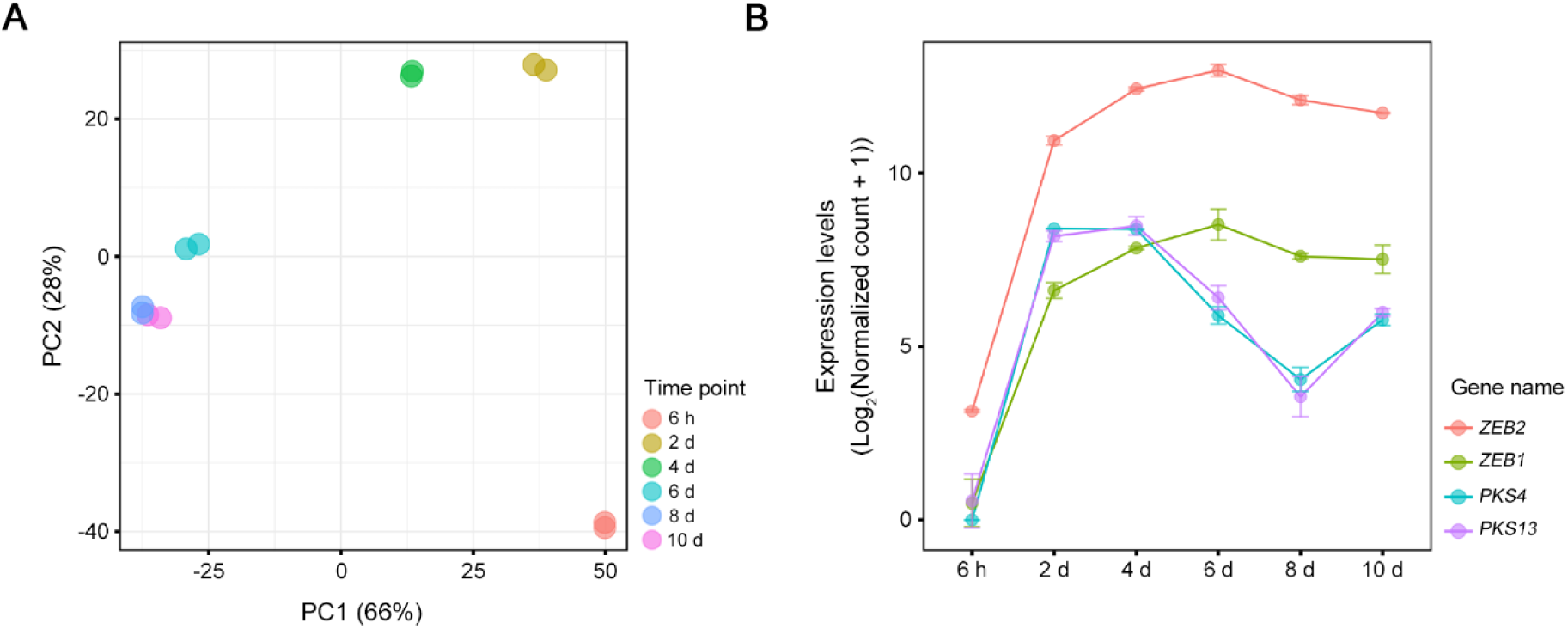
Global transcriptional trends and expression profiles of zearalenone biosynthetic genes. (A) Principal component analysis (PCA) of total transcriptome profiles. The PCA plot displays the distribution of RNA-seq samples based on total gene expression data at 6 time points (6 h, 2 d, 4 d, 6 d, 8 d, and 10 d) during zearalenone induction. Colors indicate different time points, and each point represents one of two biological replicates performed per time point. (B) Temporal expression patterns of four representative biosynthetic genes. Line graphs show the log2-transformed normalized expression levels for *ZEB2*, *ZEB1*, *PKS4*, and *PKS13*. The x-axis represents the time course of induction, and the y-axis represents the expression levels. Points and error bars indicate the mean and standard deviation of two biological replicates, respectively.

The expression patterns of core ZEA biosynthetic genes were examined in parallel with these transcriptomic changes (Fig. 1B). *ZEB2*, *ZEB1*, *PKS4*, and *PKS13* were strongly induced at 2 d compared with the 6 h time point. *ZEB1* and *ZEB2* remained highly expressed at subsequent time points, whereas *PKS4* and *PKS13* reached their highest expression levels at 2 d and gradually declined thereafter. Thus, the major transcriptomic transition between 6 h and 2 d coincided with the induction of the ZEA biosynthetic gene cluster.

Genome-wide differential expression analysis identified 3,570 differentially expressed genes (DEGs) at 2 d, 4,225 at 4 d, 5,312 at 6 d, 5,646 at 8 d, and 5,633 at 10 d relative to the 6 h time point. Across these comparisons, 7,569 unique DEGs were identified and subjected to temporal expression profiling. A hierarchical clustering heatmap revealed distinct temporal expression patterns consistent with the PCA results, with the 2 d and 4 d samples clustering together, the 6 d, 8 d, and 10 d samples forming a second cluster, and the 6 h samples exhibiting a distinct expression profile (Fig. 2A).

**Fig. 2.**
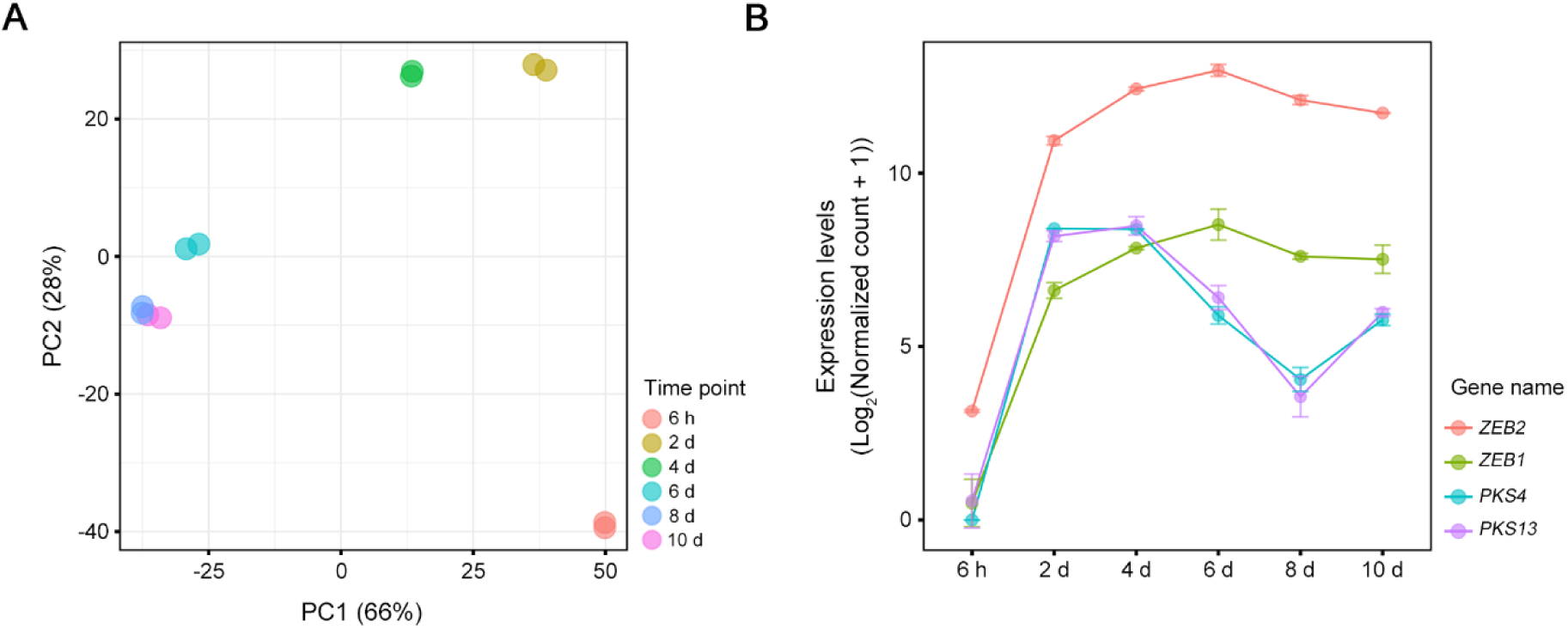
Differentially expressed genes (DEGs) under zearalenone-induction conditions. (A) Hierarchical clustering of differentially expressed genes. The heatmap displays the expression profiles of DEGs identified by comparing each subsequent time point (2 d, 4 d, 6 d, 8 d, and 10 d) to the initial 6 h reference point. The rows present individual genes, and the columns represent biological replicates for each time point. Relative expression levels are indicated by color intensity, based on VST-normalized counts scaled by z-score across rows. (B) Mfuzz clustering diagram of DEGs. The total set of DEGs was classified into six distinct clusters based on their temporal expression trajectories. The x-axis represents the time course of induction, and the y-axis indicates the standardized expression values within each cluster. Each line represents a single gene, with colors reflecting the membership values of the genes to their respective clusters.

Mfuzz analysis was performed to group genes with similar time-dependent expression patterns, which resolved the DEGs into six temporal expression clusters (Fig. 2B). Cluster 1 showed a marked decrease in expression at 2 d followed by sustained low expression. Clusters 2 and 5 were rapidly induced at 2 d but differed in their subsequent expression patterns, with cluster 2 declining markedly at later time points and cluster 5 showing a more gradual decrease. Cluster 3 exhibited a progressive decline throughout cultivation, whereas cluster 4 showed increasing expression during the early transition and continued to increase until 6 d. Cluster 6 remained relatively stable during the early stages and increased after 4 d.

Functional enrichment analysis revealed distinct biological characteristics among the temporal clusters (Fig. 3). Gene Ontology (GO) analysis showed that cluster 1 was strongly enriched for translation-associated processes, including ribosome biogenesis, rRNA processing, regulation of translation, and formation of the cytoplasmic translation-initiation complex, indicating coordinated regulation of the cellular protein-synthesis machinery (Fig. 3A). Consistent with this pattern, ribosome-related pathways were also significantly enriched in the KEGG analysis (Fig. 3B). Cluster 3 showed a similar functional pattern in both GO and KEGG analyses and was predominantly associated with central metabolic activity, including amino acid biosynthesis and metabolism, organic acid metabolism, oxidative phosphorylation, carbon metabolism, propanoate metabolism, and 2-oxocarboxylic acid metabolism, suggesting coordinated changes in nutrient utilization and energy production. Cluster 4 was enriched for DNA-templated transcription and DNA-binding-related functions in the GO analysis, whereas no significant KEGG pathway enrichment was detected. Cluster 6 also showed consistent enrichment across GO and KEGG analyses for carbohydrate metabolism and degradation, including carbohydrate and polysaccharide catabolic processes, xylan catabolism, starch and sucrose metabolism, and other glycan-degradation pathways.

**Fig. 3.**
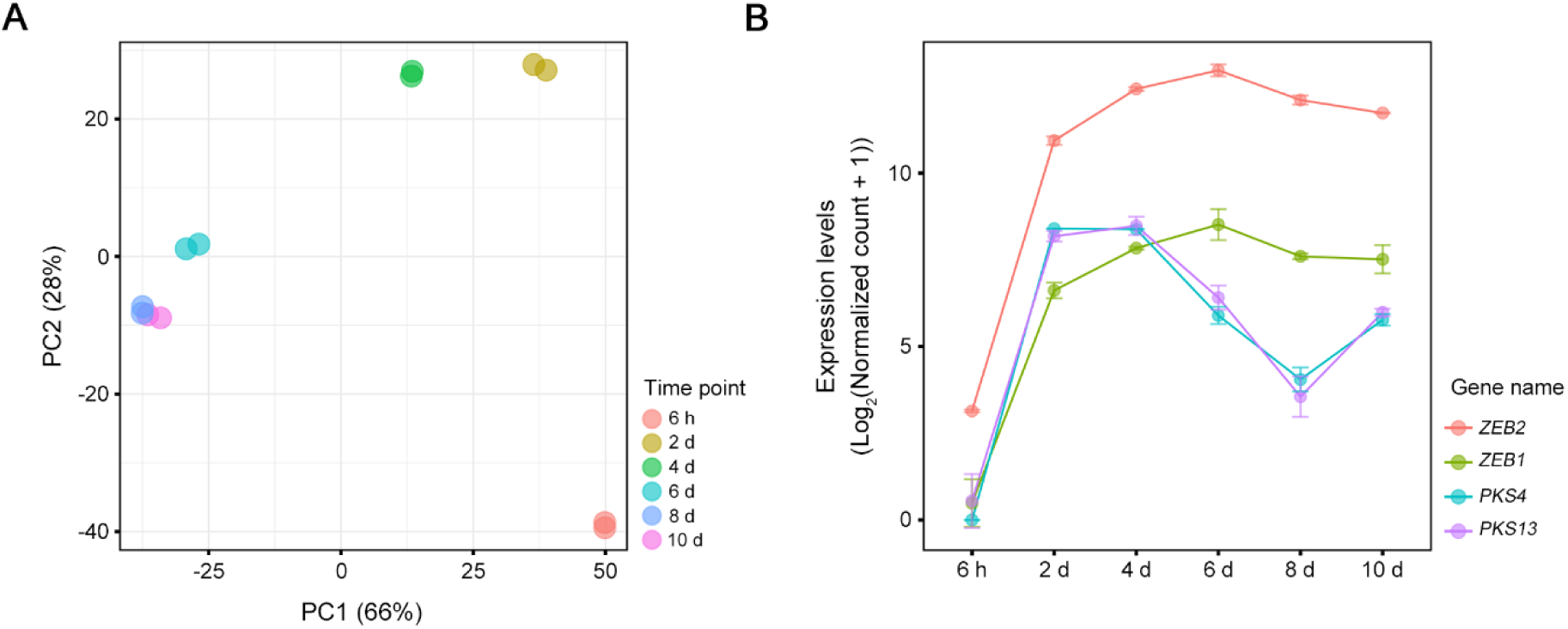
Functional enrichment analysis of DEGs within each cluster. (A) Gene Ontology (GO) enrichment analysis of cluster-specific genes. The dot plot displays the top five significant terms in the biological process category for each of the six clusters. The x-axis indicates the clusters, and the y-axis represents the enriched GO terms. The color of the dots represents the adjusted p-values, with a significant threshold of 0.05, and the size of the dots corresponds to the gene count assigned to each term. (B) Kyoto Encyclopedia of Genes and Genomes (KEGG) pathway enrichment analysis of cluster-specific genes. The dot plot shows the top five significant KEGG pathways identified within each cluster. The x-axis indicates the clusters, and the y-axis represents the pathway names. The dot color indicates the adjusted *p*-values, and the dot size reflects the gene count associated with each pathway.

Although clusters 2 and 5 did not show significant enrichment for GO biological-process terms, KEGG analysis revealed distinct pathway-level signatures. Cluster 2 was enriched for glutathione metabolism and ABC transporters, whereas cluster 5 was associated with ABC transporters, fatty acid biosynthesis, and autophagy, suggesting coordinated regulation of membrane-associated metabolism, transport, and cellular recycling processes. Collectively, these results indicate that the temporal clusters captured distinct transcriptional programs involving protein synthesis, primary metabolism, transcriptional regulation, carbohydrate metabolism, stress-associated metabolism, and intracellular transport.

### 3.2. Identification of candidate Zeb2-responsive genes associated with ZEA production

To define the putative Zeb2-responsive genes, clusters containing the core ZEA biosynthetic genes were prioritized. *ZEB2* and *ZEB1* were assigned to cluster 5, whereas *PKS4* and *PKS13* were assigned to cluster 2. Cluster 4 was also included because genes in this cluster showed increased expression during the early transition from 6 h to 2 d, followed by a further increase until 6 d. In total, 3,329 genes from clusters 2, 4, and 5 were subjected to further screening for Zeb2-dependent expression.

To identify Zeb2-responsive genes among these candidates, we performed comparative transcriptome analyses of the *ZEB2* deletion (Δ*zeb2*) and overexpression (*ZEB2OE*) strains relative to the wild-type strain. ZEA production in the wild-type strain was monitored at 6 h, 2 d, 4 d, and 6 d, whereas the Δ*zeb2* and *ZEB2OE* strains were examined at 6 h, 2 d, and 4 d. ZEA was first detected in the wild-type strain at 4 d (Fig. 4A), whereas no ZEA was detected in Δ*zeb2* during the examined period. In contrast, *ZEB2OE* showed markedly elevated ZEA production as early as 6 h (Fig. 4B). Based on these profiles, 2 d was selected for comparison between the wild type and Δ*zeb2* to capture transcriptional changes preceding detectable ZEA accumulation, whereas 6 h was selected for comparison between the wild type and *ZEB2OE* to examine early transcriptional responses associated with *ZEB2* overexpression.

**Fig. 4.**
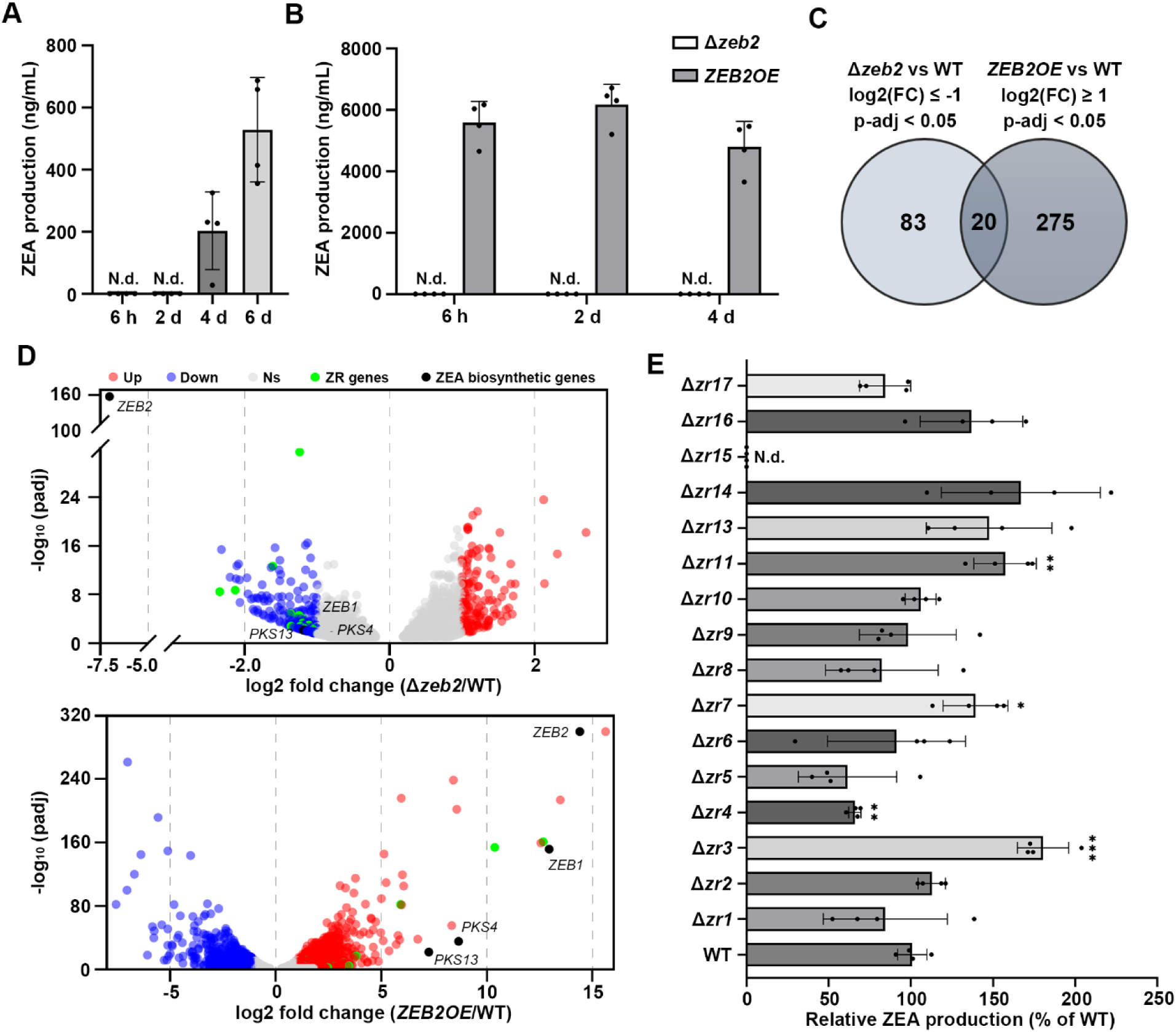
Identification of Zeb2-responsive genes associated with ZEA production. (A) ZEA production of wild-type strain cultured in SG medium for 6 h, 2 d, 4 d, and 6 d. (B) ZEA production of Δ*zeb2* and *ZEB2OE* strains at 6 h, 2 d, and 4 d. (C) Venn diagram showing the overlap between genes with reduced expression in Δ*zeb2* relative to wild type at 2 d [log2(FC) ≤ -1, adjusted p < 0.05] and genes with increased expression in *ZEB2OE* relative to wild type at 6 h [log2(FC) ≥ 1, adjusted p < 0.05]. Twenty genes, including ZEA biosynthetic cluster genes, showed overlapping Zeb2-responsive expression patterns. (D) Volcano plots showing differential gene expression in Δ*zeb2* at 2 d (upper) and *ZEB2OE* at 6 h (lower) relative to wild type. Upregulated, downregulated, and nonsignificant genes are shown in red, blue, and grey, respectively. ZR genes are shown in green, and the core ZEA biosynthetic genes are shown in black. (E) Relative ZEA production of the ZR deletion mutants. ZEA production was normalized to that of wild type, which was set to 100%. Bars represent the mean ± SD, and dots indicate individual biological replicates. Asterisks indicate significant differences compared with the wild type (\**p* < 0.05, \*\**p* < 0.01, \*\*\**p* < 0.001). N.d., not detected.

RNA-seq analysis at these time points identified 103 genes within clusters 2, 4, and 5 that showed reduced expression in Δ*zeb2* relative to the wild type at 2 d. Conversely, 295 genes within these clusters showed increased expression in *ZEB2OE* relative to the wild type at 6 h. Based on these expression profiles, 20 genes showing reciprocal expression patterns, with reduced expression in Δ*zeb2* and increased expression in *ZEB2OE*, were selected for further analysis (Fig. 4C). Volcano plot analysis illustrated the broader transcriptional responses to *ZEB2* deletion and overexpression (Fig. 4D). The core ZEA biosynthetic genes, including *PKS4*, *PKS13*, and *ZEB1*, were downregulated in Δ*zeb2* and upregulated in *ZEB2OE*, and the overlapping candidate genes exhibited the same directional response. The expected reciprocal responses of the known ZEA biosynthetic genes supported the use of this expression pattern to identify additional Zeb2-responsive candidates. After excluding the three previously characterized ZEA biosynthetic genes present among the 20 candidates (*PKS4*, *PKS13*, and *ZEB1*), the remaining 17 genes were designated as candidate Zeb2-responsive (ZR) genes (Table 1). Among the candidate ZR genes, six encoded hypothetical or uncharacterized proteins, whereas the annotated candidates included proteins putatively involved in redox metabolism, detoxification, transport, signaling, and secondary metabolism.

**Table 1.** Candidate Zeb2-responsive genes identified by comparative transcriptome analysis.

| ZR genes | Gene ID | Description | ZEB2OE/WT<br>log2(FC) | Δzeb2/WT<br>log2(FC) | Cluster |
| --- | --- | --- | --- | --- | --- |
| ZR1 | FGRAMPH1_01G12233 | nadph2:quinone reductase | 5.9 | -1.2 | 2 |
| ZR2 | FGRAMPH1_01G09373 | nonribosomal peptide synthetase | 3.7 | -1 | 5 |
| ZR3 | FGRAMPH1_01G21921 | hypothetical protein | 3.2 | -2.1 | 2 |
| ZR4 | FGRAMPH1_01G04309 | 2'-hydroxyisoflavone reductase | 2.6 | -1.1 | 2 |
| ZR5 | FGRAMPH1_01G09071 | 2og-fe oxygenase family | 1.4 | -2.3 | 2 |
| ZR6 | FGRAMPH1_01G12823 | amino-acid permease inda1 | 1.4 | -1.2 | 2 |
| ZR7 | FGRAMPH1_01G01403 | hypothetical protein | 1.4 | -1.2 | 2 |
| ZR8 | FGRAMPH1_01G13225 | hypothetical protein | 1.3 | -1 | 2 |
| ZR9 | FGRAMPH1_01G04289 | microsomal epoxide hydrolase | 1.1 | -1 | 2 |
| ZR10 | FGRAMPH1_01G00471 | glutathione s-transferase | 1 | -1 | 2 |
| ZR11 | FGRAMPH1_01G05757 | serine/threonine-protein kinase 6 | 10.3 | -1.6 | 4 |
| ZR12 | FGRAMPH1_01G08411 | hypothetical protein | 3.2 | -1.1 | 4 |
| ZR13 | FGRAMPH1_01G08413 | hypothetical protein | 1.7 | -1.2 | 4 |
| ZR14 | FGRAMPH1_01G04427 | hypothetical protein | 1.6 | -1.2 | 4 |
| ZR15 | FGRAMPH1_01G05745 | nonribosomal peptide synthetase | 11.9 | -1.2 | 5 |
| ZR16 | FGRAMPH1_01G16279 | ankyrin-1 isoform x6 | 2.1 | -1 | 5 |
| ZR17 | FGRAMPH1_01G07279 | benomyl methotrexate resistance | 1.4 | -1.3 | 5 |

Individual deletion mutants were generated to assess the contribution of these candidates to ZEA production. Targeted gene deletion was confirmed by Southern hybridization, and mutants were successfully obtained for all candidates except *ZR12* (Fig. S1). Most ZR deletion mutants showed no reduction in ZEA production, and several produced higher levels of ZEA than the wild-type strain (Fig. 4E). In contrast, ZEA was not detected in the *ZR15* deletion mutant, suggesting that *ZR15* contributes to ZEA biosynthesis.

### 3.3. *NRPS15* contributes to ZEA production in *F. graminearum*

Sequence analysis revealed that *ZR15* encodes non-ribosomal peptide synthetase 15 (*NRPS15*), which is located adjacent to the ZEA biosynthetic gene cluster (Fig. 5A). To investigate the functional role of *NRPS15*, we generated *NRPS15*-complemented and overexpression strains and examined their phenotypes. Although deletion of *NRPS15* resulted in undetectable ZEA production, introduction of the wild-type *NRPS15* allele into Δ*nrps15* restored ZEA production to wild-type levels, whereas the *NRPS15* overexpression strain produced ZEA at levels comparable to those of the wild-type strain (Fig. 5B). These results confirmed that the reduction in ZEA production was associated with loss of *NRPS15*.

**Fig. 5.**
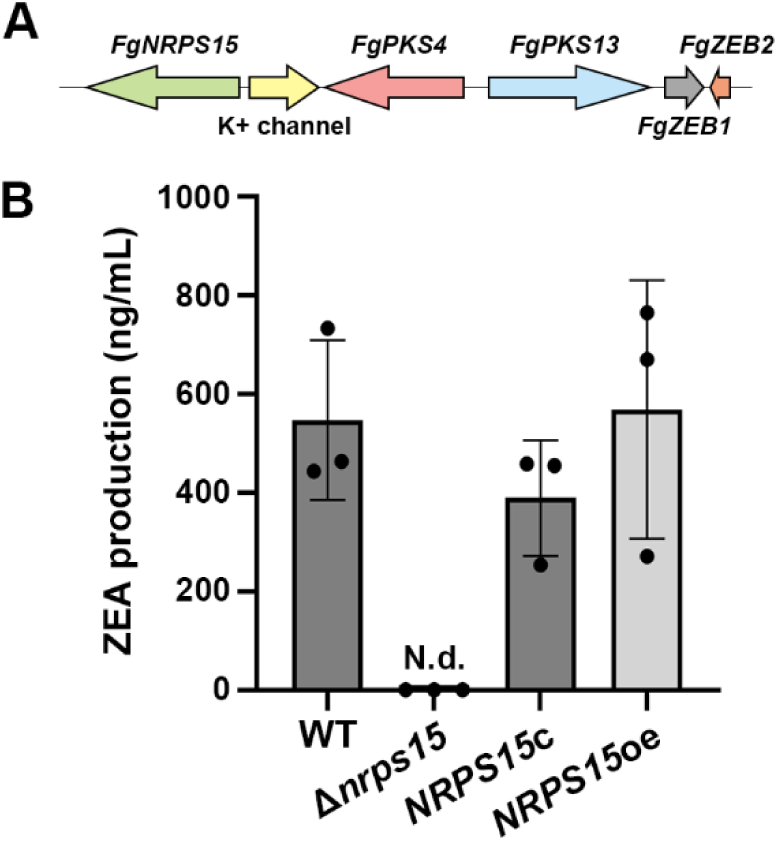
*NRPS15* contributes to ZEA production in *F. graminearum*. (A) Schematic representation of the genomic region containing *NRPS15* and the canonical ZEA biosynthetic genes *PKS4*, *PKS13*, *ZEB1*, and *ZEB2*. The neighboring K+ channel-encoding gene is also indicated. Arrows denote gene orientation. (B) ZEA production by the wild-type, Δ*nrps15*, complemented strain, and overexpressed strain after inoculation in SG medium. Bar represents the mean ± SD, and dots indicate individual biological replicates. N.d., not detected.

To determine whether *NRPS15* contributes to other biological processes, we assessed vegetative growth, conidiation, sexual development, and virulence in the mutant strains (Fig. S2A-D). The Δ*nrps15* strain showed no detectable defects in vegetative growth, conidiation, sexual development, or virulence compared with the wild-type strain.

### 3.4. *NRPS15* is associated with the expression of the ZEA biosynthetic gene cluster

Expression analysis of the core ZEA biosynthetic genes was performed to assess the effect of *NRPS15* deletion on ZEA cluster gene expression. Transcript levels of *PKS4*, *PKS13*, *ZEB1*, and *ZEB2* were significantly reduced in Δ*nrps15* compared with the wild-type strain (Fig. 6A-D). These results indicate that deletion of *NRPS15* is associated with reduced transcript levels of the ZEA biosynthetic genes.

**Fig. 6.**
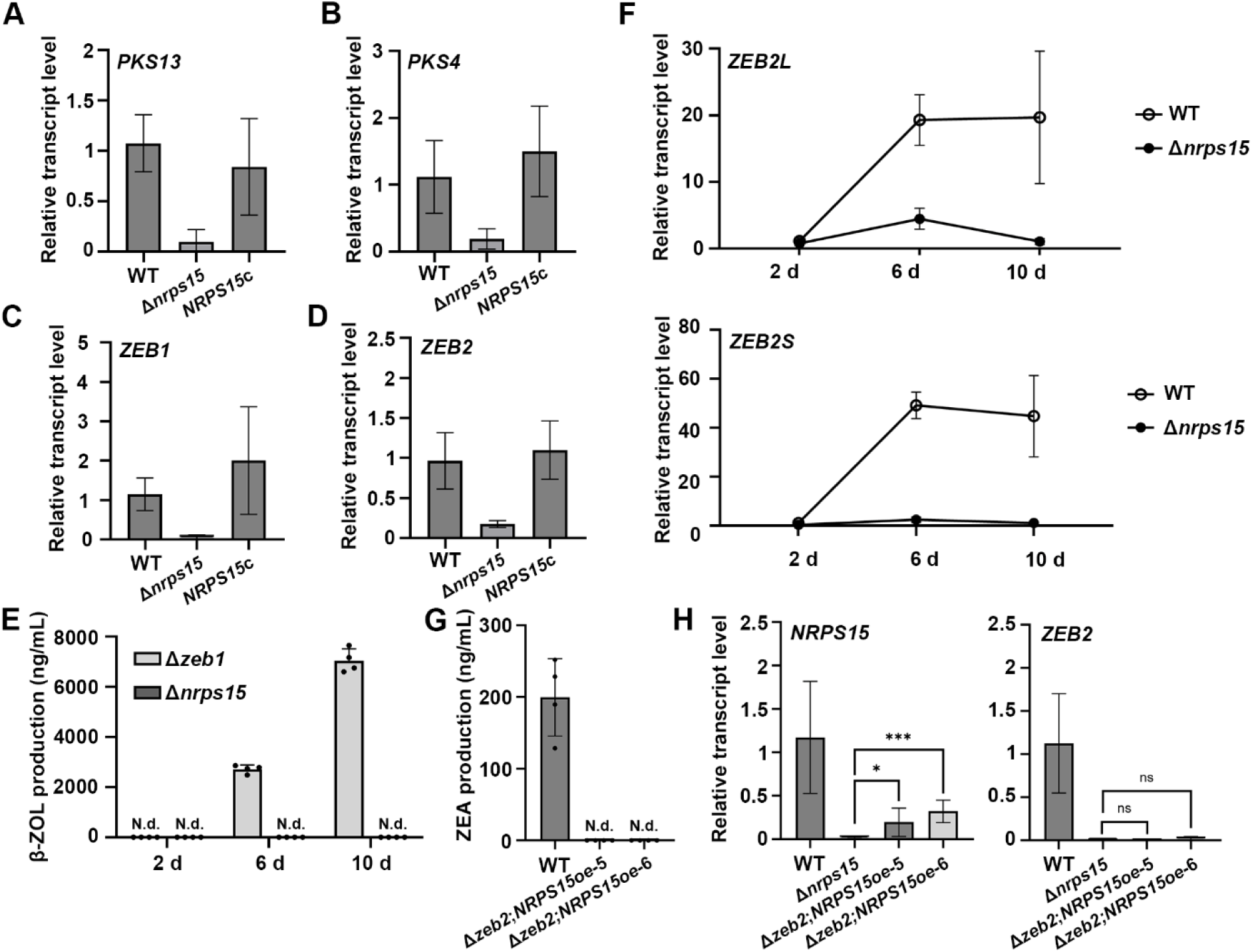
*NRPS15* is associated with expression of ZEA biosynthetic gene cluster and Zeb2-dependent ZEA production. Relative transcript levels of *PKS13* (A), *PKS4* (B), *ZEB1* (C), and *ZEB2* (D) compared to the wild type. (E) β-Zearalenol (β-ZOL) production by Δ*zeb1* and Δ*nrps15* after cultivation in SG medium for 2, 6, and 10 days. (F) Relative transcript levels of the *ZEB2L* and *ZEB2S* isoforms in wild type and Δ*nrps15* during cultivation in SG medium for 2, 6, and 10 days. (G) ZEA production by wild-type and Δ*zeb2* strains expressing *NRPS15*. (H) Relative transcript levels of *NRPS15* and *ZEB2* in wild type, Δ*nrps15*, Δ*zeb2*; *NRPS15*oe-5, and Δ*zeb2*; *NRPS15*oe-6. Asterisks indicate significant differences between the indicated strains (\**p* < 0.05, \*\*\**p* < 0.001); ns, not significant. N.d., not detected.

The accumulation of β-zearalenol (β-ZOL) was further examined to assess whether deletion of *NRPS15* altered the accumulation of ZEA-related metabolites. β-ZOL accumulated in the Δ*zeb1* mutant, whereas neither ZEA nor β-ZOL was detected in Δ*nrps15* (Fig. 6E). Together with the reduced expression of the ZEA biosynthetic genes, the absence of β-ZOL suggests that disruption of *NRPS15* affects ZEA pathway activity before substantial accumulation of ZEA or β-ZOL.

Alternative promoter usage of *ZEB2* generates two transcript isoforms, *ZEB2L* and *ZEB2S*, with opposing effects on ZEA production. Transcript levels of both *ZEB2L* and *ZEB2S* were markedly reduced in Δ*nrps15* compared with the wild-type strain (Fig. 6F), demonstrating that deletion of *NRPS15* markedly reduces the abundance of both *ZEB2* transcripts.

The functional relationship between *NRPS15* and *ZEB2* was further examined by expressing *NRPS15* in the Δ*zeb2* background. *NRPS15* expression failed to restore ZEA production in the absence of *ZEB2* (Fig. 6G). Expression of *NRPS15* in the Δ*zeb2*; *NRPS15*oe strains and the absence of *ZEB2* transcripts were confirmed by RT-qPCR (Fig. 6H). Collectively, these results indicate that the effect of *NRPS15* on ZEA production is dependent on the presence of *ZEB2* under the conditions tested.

### 3.5. Heterologous expression of *NRPS15* does not reveal a specific metabolite

Conserved domain and antiSMASH analyses revealed an NRPS-like multidomain architecture comprising A₁–CP₁–C–A₂–CP₂–R, with the C-terminal region assigned as a putative NAD(P)H-dependent thioester reductase domain (Fig. S3A). PARAS predicted serine as the preferred substrate of the first adenylation domain and alanine or proline as candidate substrates of the second. Based on these predictions, Ser–Ala- and Ser–Pro-derived products were considered candidate metabolites for targeted analysis.

To investigate whether *NRPS15* expression resulted in the production of a detectable metabolite, *NRPS15* was cloned into the pYES2 vector and heterologously expressed in *S. cerevisiae*. RT-qPCR confirmed an approximately 1.8 × 10^6^-fold increase in *NRPS15* transcript abundance in the pYES2-FgNRPS15 strain relative to the empty-vector control (Fig. S3B). Culture filtrates and cell extracts from the *NRPS15*-expressing strain were then compared with those from the empty-vector control using multiple analytical approaches. HILIC–HPLC analysis revealed broadly similar chromatographic profiles between the two strains, with no reproducible peaks specifically associated with *NRPS15* expression in either sample fraction (Fig. S4A). Consistently, UPLC–ESI–MS analysis in positive-ion mode showed no chromatographic peaks that were unique to or consistently enriched in the *NRPS15*-expressing strain (Fig. S4B). Targeted selected-ion monitoring (SIM) was subsequently performed for the predicted protonated ions of the putative Ser–Ala-derived aldehyde, alcohol, and pyrazinone at m/z 161, 163, and 141, respectively, and the corresponding Ser–Pro-derived products at m/z 187, 189, and 167; however, none of these ions showed a reproducible signal specifically associated with *NRPS15* expression (Fig. S5A). Comparison of composite positive-ion full-scan mass spectra generated from spectra acquired between 0.01 and 7.00 min revealed no distinct ions associated with *NRPS15* expression in either the culture filtrates or cell extracts (Fig. S5B). To broaden metabolite recovery, culture filtrates were sequentially extracted with EtOAc and *n*-BuOH, and RP-HPLC analysis of both solvent fractions revealed no reproducible *NRPS15*-specific peaks (Fig. S6A). The solvent extracts were subsequently subjected to trimethylsilylation and analyzed by GC–MS, but no peaks unique to or consistently enriched in the *NRPS15*-expressing strain were detected (Fig. S6B). Collectively, no reproducible metabolite specifically associated with *NRPS15* expression was detected under the cultivation, extraction, and analytical conditions employed.

### 3.6. *NRPS15* is conserved and contributes to ZEA production in *F. culmorum*

Sequence comparison revealed that *NRPS15* is conserved among several *Fusarium* species containing orthologs of the canonical ZEA biosynthetic genes, including *F. culmorum*, *F. pseudograminearum*, and *F. sporotrichioides* (Fig. 7A). The *F. culmorum* and *F. pseudograminearum* Nrps15 orthologs shared 91.1% and 93.2% amino acid identity, respectively, with *F. graminearum NRPS15*, whereas the *F. sporotrichioides* homolog showed approximately 85% identity. More distantly related sequences identified in other *Fusarium* species showed less than 40% amino acid identity and were not included in further comparative analyses.

**Fig. 7.**
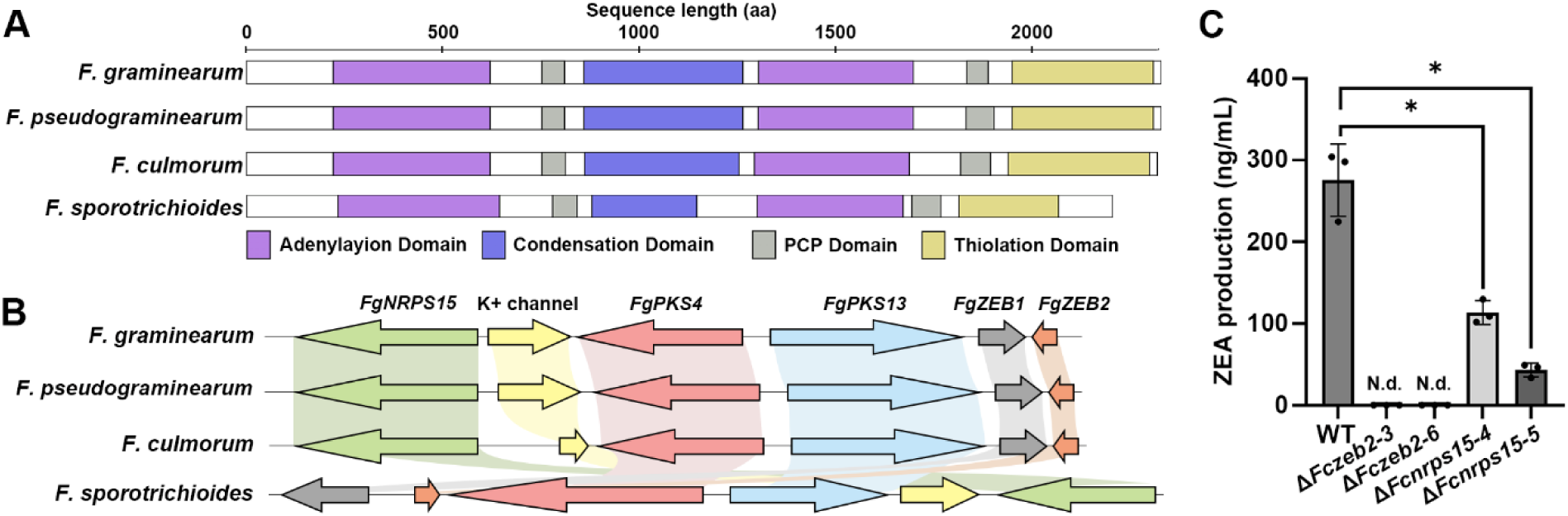
Conservation and functional contribution of *NRPS15* to ZEA production in *Fusarium* species. (A) Comparison of the predicted *NRPS15* domain architectures in *F. graminearum*, *F. pseudograminearum*, *F. culmorum*, and *F. sporotrichioides*. Conserved adenylation, condensation, PCP, and C-terminal domains are indicated according to the domain predictions. (B) Comparison of the genomic regions surrounding *NRPS15* and the ZEA biosynthetic gene cluster in the four *Fusarium* species. Homologous genes are indicated by the same colors, and connecting regions indicate sequence conservation between species. (C) ZEA production by the wild type, Δ*zeb2* mutants, and Δ*nrps15* mutants of *F. culmorum*. Bar represents the mean ± SD, and dots indicate individual biological replicates. N.d., not detected. Asterisks indicate significant differences compared to the wild type (\**p* < 0.05).

CDD analysis showed that the highly conserved *NRPS15* orthologs shared conserved NRPS domain architectures. Specifically, in *F. culmorum* and *F. pseudograminearum*, whose *NRPS15* orthologs showed >90% amino acid identity to *F. graminearum NRPS15*, the ZEA biosynthetic gene cluster was conserved and *NRPS15* was located at the corresponding genomic locus (Fig. 7B). In contrast, although the *F. sporotrichioides* homolog retained relatively high sequence identity (∼85%), the ZEA biosynthetic genes and *NRPS15* were dispersed rather than maintained in a conserved cluster arrangement.

To determine whether the contribution of *NRPS15* to ZEA production is conserved beyond *F. graminearum*, *ZEB2* and *NRPS15* deletion mutants were generated in *F. culmorum*. Similar to the phenotype observed in *F. graminearum,* deletion of *ZEB2* markedly reduced ZEA production in *F. culmorum* compared with the wild-type strain (Fig. 7C). Notably, deletion of *NRPS15* also resulted in a substantial reduction in ZEA production. These results demonstrate that *NRPS15* contributes to ZEA production in both *F. graminearum* and *F. culmorum* and provide evidence that this function is conserved at least between these two ZEA-producing *Fusarium* species.

## 4. Discussion

Transcriptome profiling combined with genetic perturbation of pathway-specific regulators has been widely used to identify previously unrecognized genes associated with fungal secondary metabolism (Nasmith et al., 2011; Price et al., 2006; Shostak et al., 2020). In particular, temporal transcriptome analysis can provide insight into regulatory events that precede the accumulation of secondary metabolites and thereby facilitate the identification of genes associated with pathway activation. Comparative analysis of deletion and overexpression mutants can further identify genes whose expression is associated with the activity of a specific regulator. Such approaches have revealed regulatory connections extending beyond canonical biosynthetic clusters in several fungal secondary metabolite pathways.

Temporal clustering showed that ZEA production was accompanied by coordinated transcriptional changes in multiple cellular processes. The most pronounced transcriptomic transition occurred between 6 h and 2 d, whereas detectable ZEA accumulation began at 4 d. Thus, transcriptional activation of the ZEA pathway preceded measurable metabolite accumulation. A similar temporal separation has been reported for trichothecene biosynthesis in *F. graminearum*, in which induction of the core TRI genes preceded toxin accumulation by approximately 2 days (Lee et al., 2014). These observations suggest that the transcriptional changes occurring between 6 h and 2 d may represent a preparatory phase involving metabolic and cellular reprogramming required for subsequent ZEA biosynthesis, whereas the changes observed from 4 d onward may partly reflect physiological responses to ZEA accumulation or to the metabolic state associated with toxin production. Clusters 2 and 5, containing *PKS4*/*PKS13* and *ZEB1*/*ZEB2*, respectively, showed relatively limited enrichment for GO biological-process terms; however, KEGG analysis identified pathways related to glutathione metabolism, ABC transporters, fatty acid biosynthesis, and autophagy. Glutathione metabolism and ABC transporters have previously been associated with oxidative and xenobiotic stress responses, multidrug resistance, and mycotoxin-related processes in *F. graminearum* (Abou Ammar et al., 2013; Lee et al., 2011; Park et al., 2024; Zheng et al., 2022), suggesting that activation of ZEA biosynthesis may be accompanied by changes in redox homeostasis, membrane transport, and cellular stress management. The limited GO enrichment of these clusters may reflect both the functional heterogeneity of the constituent genes and the incomplete annotation of genes involved in fungal secondary metabolism. Consistent with this interpretation, more than half of the putative Tri6-associated genes identified during the analysis of trichothecene regulation in *F. graminearum* were functionally unclassified (Nasmith et al., 2011). Together, these results indicate that ZEA biosynthesis is activated within the context of a broader metabolic, stress-responsive, and transcriptional transition rather than through isolated induction of the canonical ZEA biosynthetic gene cluster.

Transcriptome analyses of pathway-specific secondary-metabolite regulators have similarly shown that their effects can extend beyond canonical biosynthetic clusters. In *Aspergillus parasiticus*, transcriptional analysis of AflR identified responsive genes outside the aflatoxin cluster (Price et al., 2006). Likewise, Tri6 in *F. graminearum* influences genes involved in primary metabolism, transport, signaling, and transcription in addition to the TRI cluster and has been linked to the regulation of another secondary metabolic pathway (Nasmith et al., 2011; Shostak et al., 2020). In the present study, *ZEB2* deletion and overexpression affected numerous genes outside the known ZEA cluster, suggesting that the transcriptional influence associated with Zeb2 may extend beyond the canonical pathway genes. However, further studies are required to determine whether these transcriptional changes result from direct or indirect regulation by Zeb2 or arise secondarily from alterations in ZEA biosynthesis.

Among the 17 candidate ZR genes, several were annotated as NADPH:quinone reductase, microsomal epoxide hydrolase, glutathione S-transferase, and a benomyl/methotrexate resistance-associated protein, suggesting a possible link between the Zeb2-responsive network and cellular detoxification or redox homeostasis (Park et al., 2024; Sevastos et al., 2017; Sollner et al., 2009; Zou et al., 2000). However, most candidate deletion mutants showed little reduction in ZEA production, and several produced more ZEA than the wild type, indicating that Zeb2-responsive expression does not necessarily reflect a direct role in ZEA biosynthesis. These genes may instead represent broader cellular responses accompanying ZEA pathway activation or secondary metabolism. In contrast, deletion of *ZR15*, which encodes *NRPS15*, resulted in loss of ZEA production, highlighting *NRPS15* as a functionally important Zeb2-responsive candidate. Consistent with this phenotype, deletion of *NRPS15* reduced the expression of *ZEB2* and other ZEA biosynthetic gene, whereas *NRPS15* overexpression failed to restore ZEA production in the absence of Zeb2. These findings suggest that *NRPS15* supports Zeb2-dependent activation of ZEA biosynthesis but cannot substitute for Zeb2 function. The reciprocal reduction of *NRPS15* expression in Δ*zeb2* and *ZEB2* expression in Δ*nrps15* suggests a functional interdependence between *NRPS15* and Zeb2. Because *NRPS15* expression is Zeb2-responsive and deletion of *NRPS15* in turn reduces *ZEB2* expression, *NRPS15* may contribute to maintaining or reinforcing Zeb2-dependent activation of the ZEA pathway. However, whether this relationship represents a regulatory feedback mechanism or an indirect consequence of altered metabolic activity remains to be determined.

The physical proximity of *NRPS15* to the canonical ZEA biosynthetic genes may also be relevant. Fungal secondary-metabolite loci frequently contain accessory genes in addition to core biosynthetic enzymes, and their functional boundaries can extend beyond initially defined clusters (Chang et al., 2009; Sieber et al., 2014). Notably, a previous genome-wide analysis of *F. graminearum* placed *NRPS15* within a larger co-regulated genomic region containing ZEA-associated genes, although its biological function was unknown (Sieber et al., 2014). In addition, *NRPS15* expression was previously reported to increase together with the canonical cluster genes under ZEA-producing conditions (Lysøe et al., 2009). Consistent with these observations, our transcriptomic analysis identified *NRPS15* as a Zeb2-responsive gene, and our genetic data demonstrated that loss of *NRPS15* reduced both ZEA production and expression of the core ZEA biosynthetic genes. Comparative genomic studies further indicated that *NRPS15* is not broadly conserved across the examined *Fusarium* species but is present in several species harboring the ZEA biosynthetic gene cluster (Hansen et al., 2015; Villani et al., 2019). Moreover, deletion of *NRPS15* in *F. culmorum* substantially reduced ZEA production, providing evidence that its contribution to ZEA production is conserved between *F. graminearum* and *F. culmorum*. Together, these findings provide functional and evolutionary support for the association of *NRPS15* with the ZEA pathway and suggest that the genomic region contributing to ZEA production may extend beyond the four canonical biosynthetic genes, although functional analysis in additional ZEA-producing species will be required to determine how broadly this relationship is conserved.

Bioinformatic analyses using antiSMASH, PARAS, and NCBI CDD supported an NRPS-like multidomain architecture for *NRPS15*, yet heterologous expression in *S. cerevisiae* did not reveal a reproducible metabolite specifically associated with *NRPS15* expression using HILIC-HPLC, UPLC-ESI-MS, RP-HPLC, or GC-MS. Yeast is widely used for heterologous reconstruction of fungal PKS and NRPS pathways because of its genetic tractability and established expression systems (Tippelt and Nett, 2021; Wang et al., 2021). Nevertheless, failure to detect a product does not exclude the possibility that Nrps15 is catalytically active. *S. cerevisiae* lacks an endogenous broad-spectrum phosphopantetheinyl transferase (PPTase) suitable for efficient activation of heterologous fungal NRPSs, and functional reconstruction often requires the introduction of a compatible PPTase (Siewers et al., 2009; Tippelt and Nett, 2021). Precursor availability and the presence of additional pathway components may also limit heterologous NRPS activity in yeast (Tippelt and Nett, 2021). Moreover, a potential product present at low abundance or with physicochemical properties not adequately captured by the analytical methods used here may escape detection (Gertsman and Barshop, 2018). Thus, further analysis in the native *Fusarium* background or in a PPTase-equipped heterologous system will be important to determine the biochemical function of Nrps15.

The failure to identify a specific Nrps15-derived metabolite leaves several possible explanations for how *NRPS15* contributes to ZEA biosynthesis. Given its conserved NRPS-like domain architecture, we initially considered the possibility that Nrps15 produces a metabolite that positively influences *ZEB2* expression or ZEA pathway activity; however, the heterologous expression analyses did not provide biochemical evidence for such a product. Alternatively, Nrps15 may contribute to ZEA production through a mechanism that is not directly dependent on the synthesis of a diffusible metabolite. Another possibility is suggested by the reciprocal expression relationship between *NRPS15* and *ZEB2*. *NRPS15* expression is Zeb2-responsive, whereas deletion of *NRPS15* reduces the abundance of *ZEB2* transcripts and other ZEA biosynthetic genes. This pattern raises the possibility that *NRPS15* participates in a regulatory reinforcement mechanism associated with *ZEB2* autoregulation, contributing to the establishment or maintenance of Zeb2-dependent pathway activity, while increased Zeb2 activity may in turn reinforce *NRPS15* expression. The genomic proximity of *NRPS15* to the canonical ZEA biosynthetic genes, together with previous evidence that neighboring genes can function as accessory components of fungal secondary-metabolite loci, is also consistent with the possibility that *NRPS15* forms part of an extended functional ZEA biosynthetic region. Nevertheless, the present data do not establish whether *NRPS15* acts upstream or downstream of Zeb2, whether the relationship between the two is direct, or whether the observed reciprocal expression reflects regulatory feedback rather than an indirect consequence of altered ZEA pathway activity. Further studies in the *F. graminearum* background will therefore be required to determine the biochemical activity of *NRPS15*, clarify its relationship with Zeb2, and establish its precise role within the functional organization of the ZEA biosynthetic pathway.

Together, our study expands the current understanding of ZEA biosynthesis by identifying *NRPS15* as a previously unrecognized genetic factor required for efficient ZEA production and associated with the expression of the canonical ZEA biosynthetic genes. These findings indicate that the functional genetic framework underlying ZEA biosynthesis extends beyond the canonical four-gene biosynthetic cluster. Elucidating the biochemical function of *NRPS15* may provide further insight into the regulatory complexity of *Fusarium* secondary metabolism and may ultimately inform future approaches aimed at reducing ZEA contamination in cereals.

## 5. Conclusion

By integrating time-resolved transcriptome profiling with comparative transcriptomic analysis of *ZEB2* deletion and overexpression strains, this study identified *NRPS15* as a previously uncharacterized Zeb2-responsive genetic factor required for efficient ZEA production in *F. graminearum*. Genetic analysis demonstrated that deletion of *NRPS15* markedly reduced ZEA production and reduced the expression of core ZEA biosynthetic genes, whereas complementation restored ZEA production to wild-type levels. The contribution of *NRPS15* to ZEA production was also observed in *F. culmorum*, providing evidence that this function is conserved between two examined ZEA-producing *Fusarium* species. These findings indicate that the functional genetic framework underlying ZEA production extends beyond the canonical four-gene biosynthetic cluster and establish *NRPS15* as an additional genetic component associated with ZEA pathway activity in *Fusarium*.

## CRediT authorship contribution statement

**Nahyun Lee:** Conceptualization, Data curation, Investigation, Methodology, Validation, Visualization, Writing – original draft. **Jae Woo Han:** Data curation, Investigation, Methodology, Visualization. **Jung-Eun Kim:** Conceptualization, Investigation, Methodology. **Jiyoung Shin:** Conceptualization, Investigation, Methodology. **Ae Ran Park:** Conceptualization, Methodology. **Yin-Won Lee:** Conceptualization, Methodology. **Hun Kim:** Data curation, Investigation. **Jiyeun Park:** Conceptualization, Validation, Visualization, Writing – original draft. **Hokyoung Son:** Conceptualization, Funding acquisition, Investigation, Methodology, Supervision, Writing – original draft.

## Declaration of Competing Interest

The authors declare that they have no known competing financial interests or personal relationships that could have appeared to influence the work reported in this paper.

## Data Availability

All RNA-seq data have been deposited in the NCBI Sequence Read Archive (SRA) under accession numbers PRJNA1521793 and PRJNA1521817.

## Acknowledgements

This study was supported by the Korea Institute of Planning and Evaluation for Technology in Food, Agriculture, and Forestry (IPET) through the Agriculture and Food Convergence Technologies Program for Research Manpower Development, funded by the Ministry of Agriculture, Food and Rural Affairs (MAFRA) (No. RS-2024-00398300) and the National Research Foundation of Korea (NRF) grants funded by the Korea government (MSIT) (No. RS-2025-00553624 and RS-2026-25555789).

## Notes

https://www.ncbi.nlm.nih.gov/bioproject/PRJNA1521793

https://www.ncbi.nlm.nih.gov/bioproject/PRJNA1521817

### Competing Interest Statement

The authors have declared no competing interest.

