## Supplementary Materials for "Transcriptome dissection of the Zeb2-associated network reveals a role for *NRPS15* in zearalenone biosynthesis in *Fusarium graminearum*"

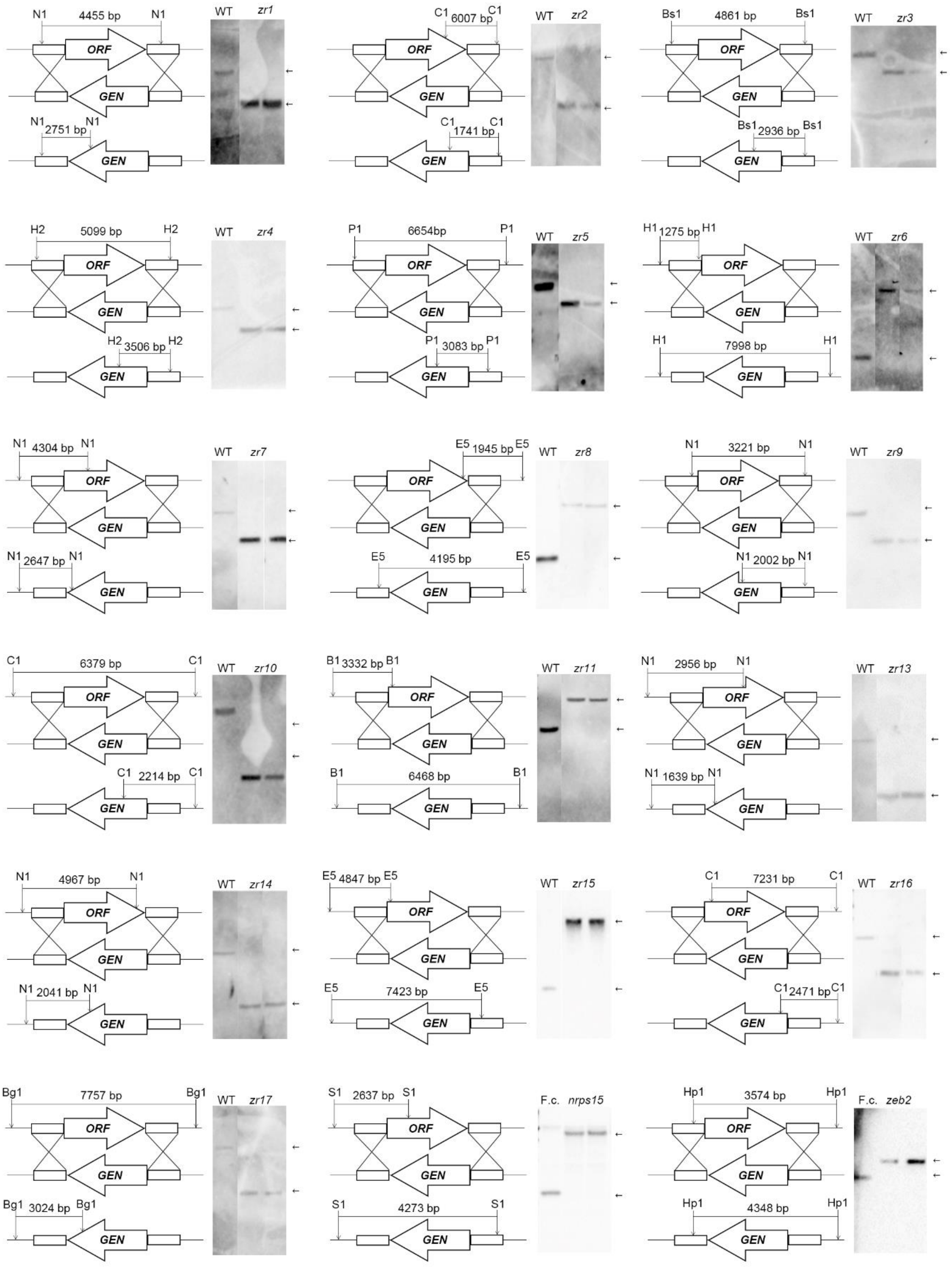

N1 4455 bp N1
ORF
GEN
N1 2751 bp N1
WT zr1
C1 6007 bp C1
C1 1741 bp C1
WT zr2
Bs1 4861 bp Bs1
Bs1 2936 bp Bs1
WT zr3
H2 5099 bp H2
H2 3506 bp H2
WT zr4
P1 6654bp P1
P1 3083 bp P1
WT zr5
H1 1275 bp H1
H1 7998 bp H1
WT zr6
N1 4304 bp N1
N1 2647 bp N1
WT zr7
E5 1945 bp E5
E5 4195 bp E5
WT zr8
N1 3221 bp N1
N1 2002 bp N1
WT zr9
C1 6379 bp C1
C1 2214 bp C1
WT zr10
B1 3332 bp B1
B1 6468 bp B1
WT zr11
N1 2956 bp N1
N1 1639 bp N1
WT zr13
N1 4967 bp N1
N1 2041 bp N1
WT zr14
E5 4847 bp E5
E5 7423 bp E5
WT zr15
C1 7231 bp C1
C1 2471 bp C1
WT zr16
Bg1 7757 bp Bg1
Bg1 3024 bp Bg1
WT zr17
S1 2637 bp S1
S1 4273 bp S1
F.c. nrps15
Hp1 3574 bp Hp1
Hp1 4348 bp Hp1
F.c. zeb2

**Fig. S1. Southern hybridization analysis of targeted gene deletion mutants.** Schematic representations of the wild-type loci and targeted gene replacement constructs are shown for the indicated ZR genes in *F. graminearum* and for *NRPS15* and *ZEB2* in *F. culmorum*. Target genes were replaced with the geneticin resistance cassette (*GEN*) by homologous recombination. Restriction sites and the expected sizes of hybridizing fragments are indicated in each schematic. Genomic DNA from the wild type and the corresponding deletion mutants was digested with the indicated restriction enzymes and subjected to Southern hybridization. Arrows indicate the expected hybridizing bands for the wild type and deletion alleles. Southern hybridization confirmed the predicted gene replacement patterns in the mutant strains. Hp1, HpaI; S1, SacI; N1, NcoI; C1, ClaI; B1, BclI; Bs1, BsaI; Bg1, BglI; H2, HaeII; H1, HgaI; E5, EcoRV; P1, PstI; *GEN*, geneticin resistance cassette; WT, wild-type strain.

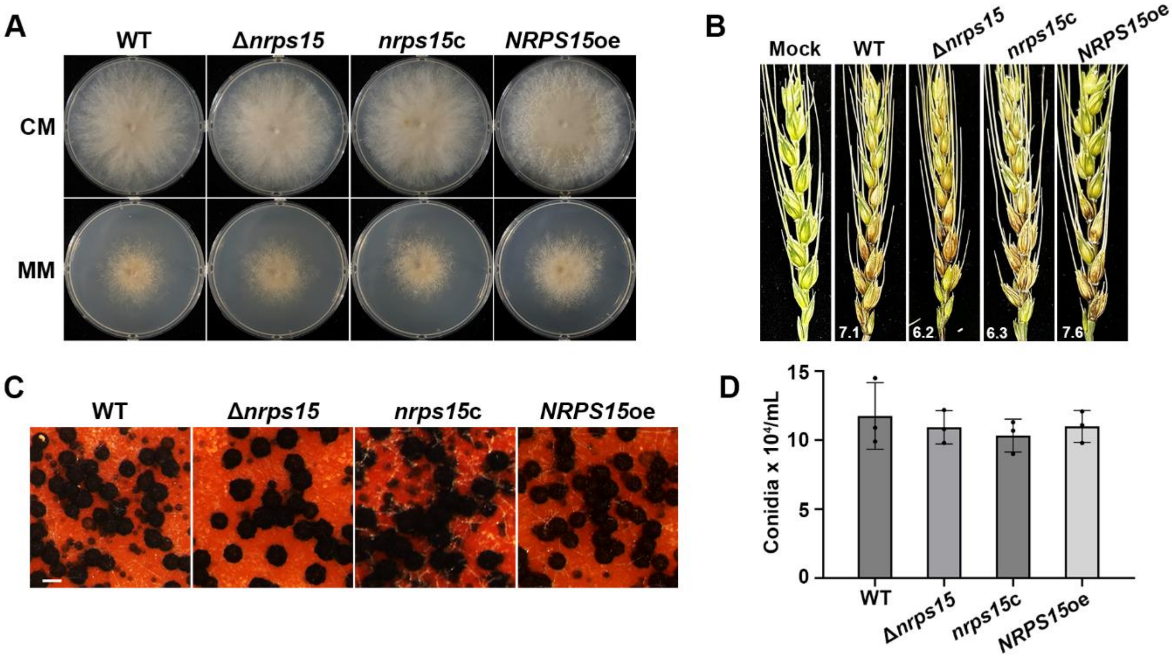


**Fig. S2. Phenotypic characterization of *NRPS15* mutant strains in *F. graminearum*.** (A) Vegetative growth of the wild-type, Δ*nrps15*, complemented, and overexpression strains on CM and MM. (B) Pathogenicity of the indicated strains on wheat heads. Mock-treated wheat heads served as negative controls, and numbers indicate the mean number of symptomatic spikelets. (C) Sexual development of the indicated strains on carrot agar. Scale bar, 200 µm. (D) Conidial production of the indicated strains after cultivation in CMC medium. Bars represent the mean ± SD, and dots indicate individual biological replicates.

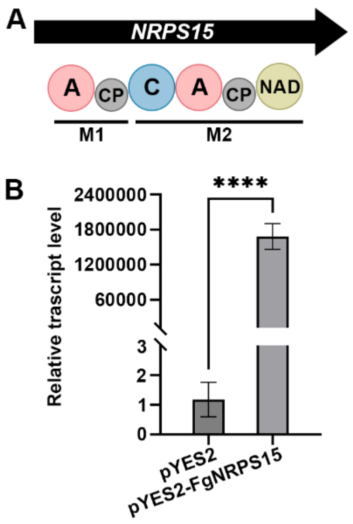


**Fig. S3. Predicted domain architecture and heterologous expression of *NRPS15* in *S. cerevisiae*.** (A) Predicted domain architecture of *NRPS15*. The protein contains two adenylation (A) domains, two carrier protein (CP) domains, a condensation (C) domain, and a C-terminal NAD(P)H-dependent reductase domain. M1 and M2 indicate the two predicted NRPS modules. (B) Relative *NRPS15* transcript levels in *S. cerevisiae* carrying the empty pYES2 vector or pYES2-FgNRPS15, determined by RT-qPCR. Bars represent the mean ± SD, and dots indicate individual biological replicates. ****$p < 0.0001$.

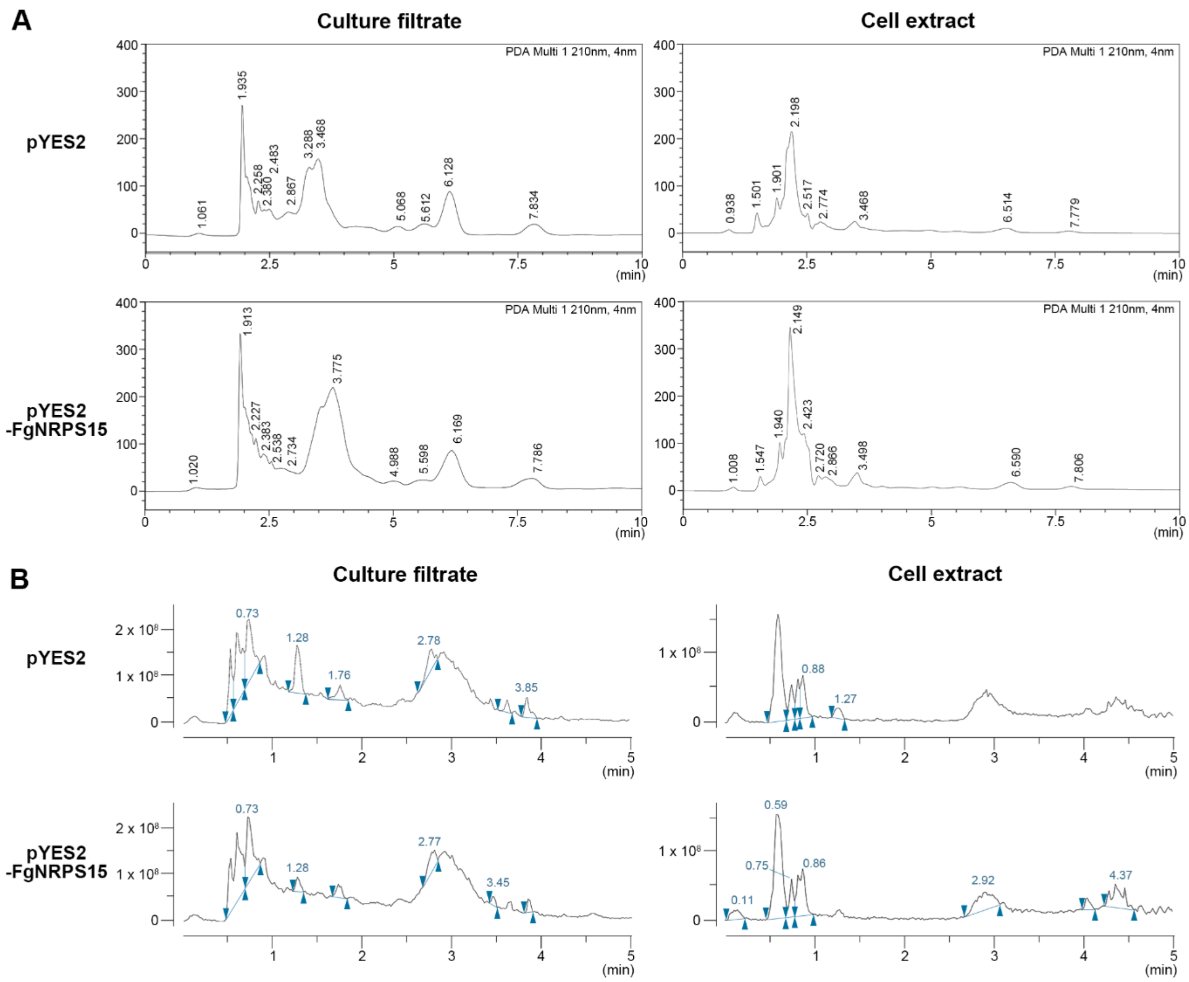


**Fig. S4. HILIC–HPLC and UPLC–ESI–MS analyses of culture filtrates and cell extracts from *NRPS15*-expressing and control yeast strains.** (A) Representative HILIC–HPLC chromatograms of culture filtrates and cell extracts from *S. cerevisiae* BY4741 carrying the empty pYES2 vector or pYES2-FgNRPS15. Samples were separated on a YMC-Triart Diol-HILIC column under isocratic conditions using acetonitrile/water (90:10, v/v) and monitored at 210 nm. (B) UPLC–ESI–MS total ion chromatograms (TICs) of culture filtrates and cell extracts from the same strains analyzed in positive-ion mode. No reproducible chromatographic peaks unique to or consistently enriched in the *NRPS15*-expressing strain were detected in either sample fraction.

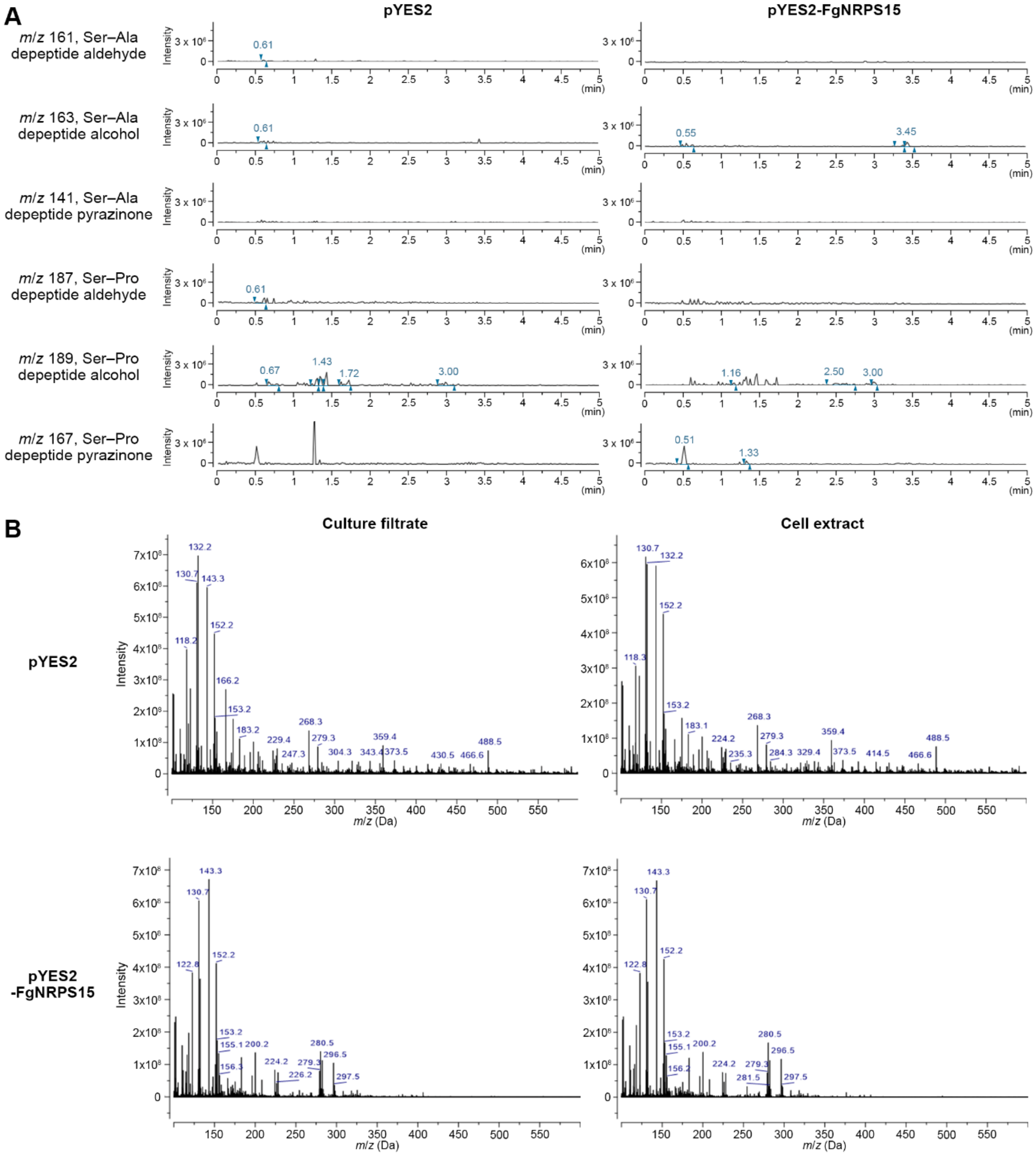


**Fig. S5. Targeted SIM and composite positive-ion mass spectral analyses of predicted *NRPS15*-derived products in *S. cerevisiae*.** (A) Selected-ion monitoring (SIM) chromatograms acquired at the predicted protonated *m*/*z* values for putative Ser–Ala-derived aldehyde, alcohol, and pyrazinone products (*m*/*z* 161, 163, and 141, respectively) and the corresponding Ser–Pro-derived products (*m*/*z* 187, 189, and 167) in *S. cerevisiae* BY4741 carrying the empty pYES2 vector or pYES2-FgNRPS15. No reproducible signals specifically associated with *NRPS15* expression were detected at the monitored ions. (B) Composite positive-ion mass spectra of culture filtrates and cell extracts from the same strains. Full-scan ESI-positive spectra acquired over the chromatographic interval from 0.01 to 7.00 min were combined to generate a composite spectrum for each sample. No distinct ions specifically associated with *NRPS15* expression were observed in either fraction.

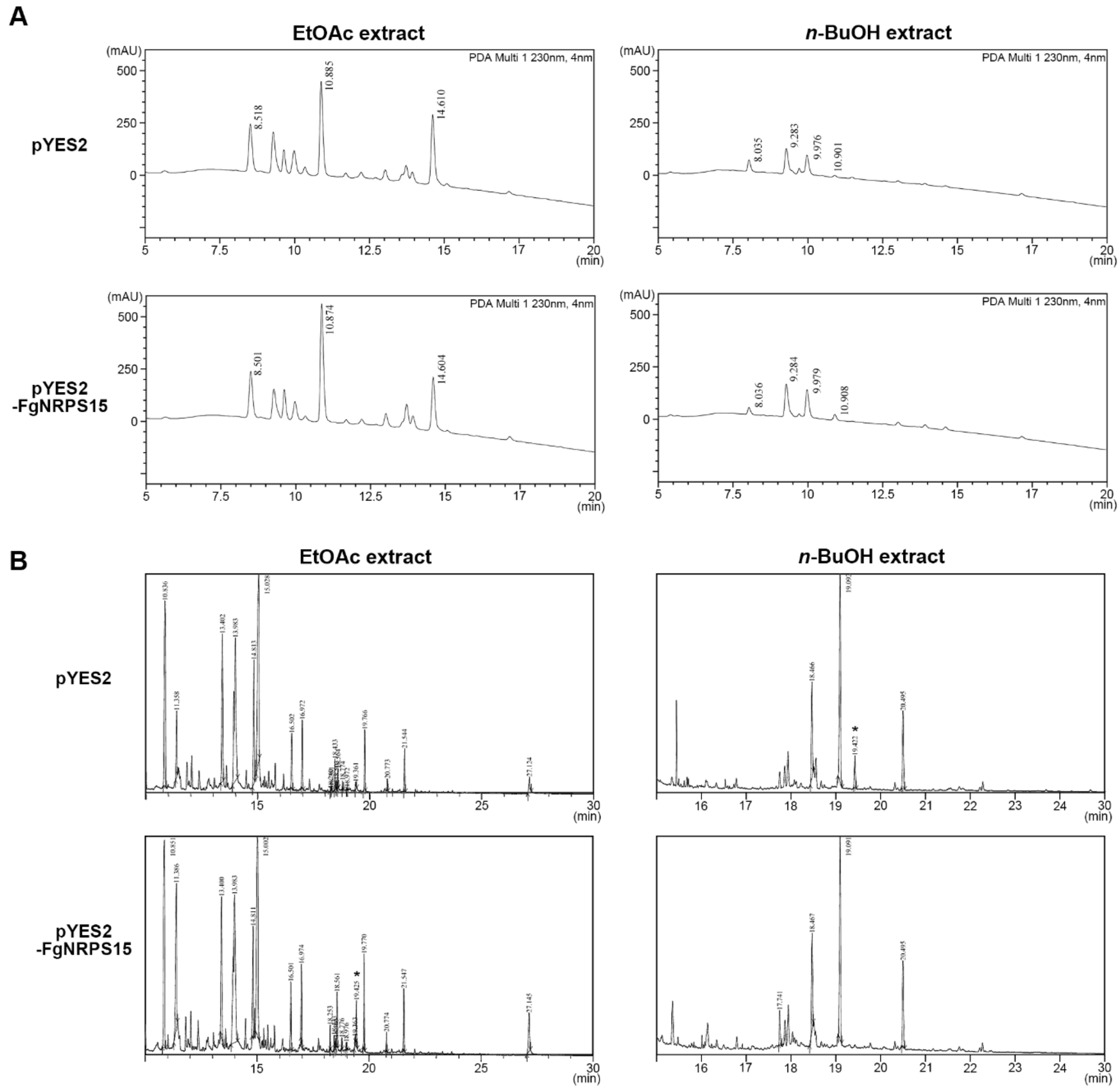


**Fig. S6. RP-HPLC and GC–MS analyses of solvent-extracted metabolites from *NRPS15*-expressing and control yeast strains.** (A) RP-HPLC chromatograms of ethyl acetate (EtOAc) and *n*-butanol (*n*-BuOH) extracts prepared from culture filtrates of *S. cerevisiae* BY4741 carrying the empty pYES2 vector or pYES2-FgNRPS15. Extracts were separated on a Phenomenex Luna C18(2) column using a linear gradient of 20–100% aqueous methanol containing 0.1% formic acid over 20 min and monitored at 230 nm. No reproducible peaks specifically associated with *NRPS15* expression were detected in either solvent fraction. (B) GC–MS total ion chromatogams of trimethylsilyl (TMS)-derivatized EtOAc and *n*-BuOH extracts from the same strains. No reproducible peaks unique to or consistently enriched in the *NRPS15*-expressing strain were detected. The asterisk indicates a peak identified as D-glucose.

**Table S1. Strains used in this study**

| Strain | Description | Reference |
|---|---|---|
| Z-3639 | *F. graminearum* wild-type | [19] |
| Δ*zeb2* | *ZEB2* deletion mutant | [18] |
| *ZEB2OE* | *ZEB2* overexpression strain | [18] |
| Δ*zr1* | *ZR1* deletion mutant | This study |
| Δ*zr2* | *ZR2* deletion mutant | This study |
| Δ*zr3* | *ZR3* deletion mutant | This study |
| Δ*zr4* | *ZR4* deletion mutant | This study |
| Δ*zr5* | *ZR5* deletion mutant | This study |
| Δ*zr6* | *ZR6* deletion mutant | This study |
| Δ*zr7* | *ZR7* deletion mutant | This study |
| Δ*zr8* | *ZR8* deletion mutant | This study |
| Δ*zr9* | *ZR9* deletion mutant | This study |
| Δ*zr10* | *ZR10* deletion mutant | This study |
| Δ*zr11* | *ZR11* deletion mutant | This study |
| Δ*zr13* | *ZR13* deletion mutant | This study |
| Δ*zr14* | *ZR14* deletion mutant | This study |
| Δ*zr15* (Δ*Fgnrps15*) | *ZR15* deletion mutant; *Fgnrps15* deletion mutant | This study |
| Δ*zr16* | *ZR16* deletion mutant | This study |
| Δ*zr17* | *ZR17* deletion mutant | This study |
| *nrps15*c | NRPS15-complemented strain | This study |
| *NRPS15*oe | NRPS15 overexpression strain | This study |
| Δ*zeb2; NRPS15*oe | Δ*zeb2* strain expressing *NRPS15* under RP27 promoter | This study |
| *Fusarium culmorum* | *F. culmorum* wild type strain | KACC 48423 |
| Δ*Fczeb2* | *F. culmorum ZEB2* deletion mutant | This study |
| Δ*Fcnrps15* | *F. culmorum NRPS15* deletion mutant | This study |
| pYES2 | *S. cerevisiae* BY4741 carrying the empty pYES2 vector | This study |
| pYES2-FgNRPS15 | *S. cerevisiae* BY4741 carrying the pYES2-FgNRPS15 vector | This study |

### Table S2. Primers used in this study

| Primer name | Sequence (5′–3′) | Purpose |
|---|---|---|
| ZR1-5F | TCGACGTCGTTGTTGGTTCTAATC | |
| ZR1-5N | CAAGTCTATTCGAGCCAGTTCACG | |
| ZR1-5R | GCACAGGTACACTTGTTTAGAGAGGAATGAGGTTTTGGACTGATG | *ZR1* deletion mutant |
| ZR1-3F | CCTTCAATATCATCTTCTGTCGTAACCAATATTCTTTAACGCCTGTCT | |
| ZR1-3N | TCAGTACTTTGAGATTTCACCTTTGG | |
| ZR1-3R | TCAAGAGAACCGAAAGGGAGATT | |
| ZR1 with 5F | AGCGATAGCCTTTGTAACGAGTCT | |
| ZR2-5F | ATGCTGGCCATTGGTAGATGC | |
| ZR2-5N | CCTACGATCATCCCCCACCTT | |
| ZR2-5R | GCACAGGTACACTTGTTTAGAGGTAGGACCAAAGGAATTGTGCTTAC | *ZR2* deletion mutant |
| ZR2-3F | CCTTCAATATCATCTTCTGTCGTACGTTAGTGCTTCACTTGGAAATG | |
| ZR2-3N | GGACGTCCCTTCGAAGTAACAGT | |
| ZR2-3R | GGAACTTTGGTACTCTTGGATGTTG | |
| ZR2 with 5F | CGTTGACGCCCCGAAGTT | |
| ZR3-5F | CAGGCGGACAGCGATTGAACT | |
| ZR3-5N | TCTGACTTGCCGAACATAAACTGA | |
| ZR3-5R | GCACAGGTACACTTGTTTAGAGGGTGTGTAGTAGCGAAGGTGTTCT | |
| ZR3-3F | CCTTCAATATCATCTTCTGTCGTCCTGGAAGCGAAGCACTTG | *ZR3* deletion mutant |
| ZR3-3N | TCATCGACACAGATCCCTCAGC | |
| ZR3-3R | TCCGACGGTACCATACGAGC | |
| ZR3 with 5F | GGTGGTGAAGCTTGGGTGAC | |
| ZR4-5F | CTCTATTCTTGCCCTCGCCAT | |
| ZR4-5N | GGCATGCCTTTACGTTGGAGAC | |
| ZR4-5R | GCACAGGTACACTTGTTTAGAGCTAATTGCCGGCTGGAACACT | |
| ZR4-3F | CCTTCAATATCATCTTCTGTCGGAGTGTTATTGCAATGGTTCTT | *ZR4* deletion mutant |
| ZR4-3N | ACACTGTGTTCCAACTTTGTCCTG | |
| ZR4-3R | GGCAGAGGTGCTAGGTTTTATCAT | |
| ZR4 with 5F | CGCCTTGACATCTTCCATTTTG | |
| ZR5-5F | TTGCAGGGTTGGCCTCATCT | |
| ZR5-5N | ACTGATAAGATGCATCTTTTTGGGAA | |
| ZR5-5R | GCACAGGTACACTTGTTTAGAGTGAGTGTTTGTTGTTGTTGTGATTG | *ZR5* deletion mutant |
| ZR5-3F | CCTTCAATATCATCTTCTGTCGTGGGTCTTTTTAGGGCTGATTC | |
| ZR5-3N | GAGCGGCCCGTTCCTGAT | |

| | | |
|---|---|---|
| ZR5-3R | GCACGGGAGCTAGAGACAGGA | |
| ZR5 with 5F | GCGCTTGCCCGAGAGGA | |
| ZR6-5F | AGTCTCGCTTCATCAAAACAAACA | *ZR6* deletion mutant |
| ZR6-5N | ACCTTGTCAATAGAGATACGCACTG | |
| ZR6-5R | GCACAGGTACACTTGTTTAGAGTCTGACAATCCAATCGA GATGTGA | |
| ZR6-3F | CCTTCAATATCATCTTCTGTCGCCTACTTGTGGTCCTTCTT TGTGT | |
| ZR6-3N | GGCTGGACCTACGTTTGGCTTAT | |
| ZR6-3R | TCCCCATGGCTTTGAACACG | |
| ZR6 with 5F | CCAAAGACGTGCACCCCAGT | |
| ZR7-5F | TCTGGGCATGGGGCAAAAC | *ZR7* deletion mutant |
| ZR7-5N | AATTGATTGATTGCATTGACTTCTG | |
| ZR7-5R | GCACAGGTACACTTGTTTAGAGATTGCGATATTTGGGAG TTGTGG | |
| ZR7-3F | CCTTCAATATCATCTTCTGTCGCGGATTTTTTTTTTGAACG GTT | |
| ZR7-3N | TCTTGGATCATCATCAGGAGACTAAT | |
| ZR7-3R | CTCTTCCCCCTCCTTGTGATAACT | |
| ZR7 with 5F | TTGGAGTACTTTGAGAGCCTGTTT | |
| ZR8-5F | GCCTCTACCATGCTTCGTCTCTTT | *ZR8* deletion mutant |
| ZR8-5N | TGTCTTCGCTTTCTCTGGTGGTTA | |
| ZR8-5R | GCACAGGTACACTTGTTTAGAGTTGTCTTGTGTTGAAGTT GGAAAG | |
| ZR8-3F | CCTTCAATATCATCTTCTGTCGCACGATGAAATGAGACA AGCACT | |
| ZR8-3N | CTCACCATCGCTTTAATCCAGTCA | |
| ZR8-3R | ATAAGGCTGGATGTTTGTCTCACT | |
| ZR8 with 5F | TGTCGCACGGCATTGTCC | |
| ZR9-5F | TTCGTCGAATCCCTCCTGG | *ZR9* deletion mutant |
| ZR9-5N | GAGCACAATACCCCTTCCTCAC | |
| ZR9-5R | GCACAGGTACACTTGTTTAGAGGATGTTGGCGTGCTAAA AAATGA | |
| ZR9-3F | CCTTCAATATCATCTTCTGTCGCATCATGGGCAGTGCAGA ATAAAT | |
| ZR9-3N | TCGCCAAGACATGTTCCCAGATA | |
| ZR9-3R | GAAATGACACCGTTCTCCACAGT | |
| ZR9 with 5F | CGATTGTGCTTGGCTTGGTG | |
| ZR10-5F | CTCTCGAGTAACTGGTGGTGACA | *ZR10* deletion mutant |
| ZR10-5N | TTTATTCCATCAGCTGCCAGTCA | |
| ZR10-5R | GCACAGGTACACTTGTTTAGAGACCTAGCATTCAGCAGC AGAGA | |
| ZR10-3F | CCTTCAATATCATCTTCTGTCGAAACGTGGCTATGCTTTG TCTACA | |
| ZR10-3N | ATAAAATGAAACGGCAAAACGATA | |
| ZR10-3R | TGAGAATGAACAGAGAGAAAAGGG | |

| | | |
|---|---|---|
| ZR10 with 5F | GCCTGCGCCCACTTGTAAA | |
| ZR11-5F | CAATGCCAACCTCGAAACACTA | *ZR11* deletion mutant |
| ZR11-5N | TCCCATAAGAGTGGGAACTGC | |
| ZR11-5R | GCACAGGTACACTTGTTTAGAGGGGTGCTGTGAATATTG AAGAAAG | |
| ZR11-3F | CCTTCAATATCATCTTCTGTCGGACAGGTGGTAGCTCCTG ATTCTA | |
| ZR11-3N | GTCTCCACCGCAACCATCTCT | |
| ZR11-3R | CTTGGTTACTCCGAGACTGAAGTT | |
| ZR11 with 5F | GACTGGCGTATCTTCAAATGGTG | |
| ZR12-5F | GCAGAGATCGGCATATTGAGGTTT | *ZR12* deletion mutant |
| ZR12-5N | AGTATGTCCCCGCCAATGAAA | |
| ZR12-5R | GCACAGGTACACTTGTTTAGAGGTAGTGATGGTAAGAAT GATGTTTGAC | |
| ZR12-3F | CCTTCAATATCATCTTCTGTCGGTTTTAGGGTATGCTTCT GATTCGG | |
| ZR12-3N | TTCGGCAAAGAGCAGTGGC | |
| ZR12-3R | TACATGGGGAGGTCACAAGCA | |
| ZR12 with 5F | ATGTCGCAGAATTGTCCGTCA | |
| ZR13-5F | CAAGTGGCATTTCTGTCGTCATTA | *ZR13* deletion mutant |
| ZR13-5N | GTTCTTGACATCTTCCCGAGTGAG | |
| ZR13-5R | GCACAGGTACACTTGTTTAGAGCCAACATGAAACCAACA CAGGATA | |
| ZR13-3F | CCTTCAATATCATCTTCTGTCGGTACCACTCGTTGTCTTTG TCCTA | |
| ZR13-3N | CAAGGATGATGGCGTGAGC | |
| ZR13-3R | GCTCGATGGATGGCAAAAG | |
| ZR13 with 5F | GGTGGTGGTGGTGAAGAAGTGAG | |
| ZR14-5F | GCGCCGATACCCTTCACG | *ZR14* deletion mutant |
| ZR14-5N | TTCCCCGGCGAGGTTGT | |
| ZR14-5R | GCACAGGTACACTTGTTTAGAGATCCGGGGTTGATATAA ATAAGCG | |
| ZR14-3F | CCTTCAATATCATCTTCTGTCGCTTTGTTTCATTGGGCTTG TGTA | |
| ZR14-3N | TGTGTGGACGGCGAGTTGT | |
| ZR14-3R | TTGCGACGCTACAACAGGTT | |
| ZR14 with 5F | TCCGATGCTCCCTGTTCTCTACTT | |
| ZR15-5F | AGCTTGATAGGAGAAAATACCAGAA | *ZR15* deletion mutant |
| ZR15-5N | CCGAGACATAATGCTTGCGAG | |
| ZR15-5R | GCACAGGTACACTTGTTTAGAGTTCTGGCCATTCAAATCG TTATC | |
| ZR15-3F | CCTTCAATATCATCTTCTGTCGAAGAGTGTGTTGTCAAAT GACTATGG | |
| ZR15-3N | ACAGGTCGGCATATCCCAAA | |
| ZR15-3R | TCGATATCCATCCCCCACAA | |
| ZR15 with 5F | TGTTTTGAGGCCCGACCATAG | |

| | | |
|---|---|---|
| ZR16-5F | ATATTGCCTCTCGGTGCCAGT | *ZR16* deletion mutant |
| ZR16-5N | TTTATCTTGCCATCCCGACACTTA | |
| ZR16-5R | GCACAGGTACACTTGTTTAGAGTGATGTTCAAGTGCCCG TGTAT | |
| ZR16-3F | CCTTCAATATCATCTTCTGTCGGATGTCAACGCGTCGGAA ATAG | |
| ZR16-3N | CAAGCCATGACCGGGATTAGATT | |
| ZR16-3R | ATCCCGACCAGGTACGAGTTC | |
| ZR16 with 5F | GGATTCGCTCCAAAGTCCAGTAG | |
| ZR17-5F | TCATCAACTCACTCGCTCAATCTG | *ZR17* deletion mutant |
| ZR17-5N | GGCCAGGGCTTGCGATAGT | |
| ZR17-5R | GCACAGGTACACTTGTTTAGAGCAAGGGTCCAAACATTT ACATACAC | |
| ZR17-3F | CCTTCAATATCATCTTCTGTCGCGCCCTACGATTCAAGAG TCAT | |
| ZR17-3N | AGGGAAGGCGATGTTTTGAGT | |
| ZR17-3R | GAGCTTCTCCTGGGTAAACACG | |
| ZR17 with 5F | GCTGAAAGACGATAGGATAGATGC | |
| FcZEB2-5F | CCTCCCGGCTGACACTCC | *FcZEB2* deletion mutant |
| FcZEB2-5N | CTCACTGCGGCTTGTAGAGTCA | |
| FcZEB2-5R | GCACAGGTACACTTGTTTAGAGTGAATTAGGGTCACAGG CGTAG | |
| FcZEB2-3F | CCTTCAATATCATCTTCTGTCGGATGATACAGCTTCCTCA CCTCTG | |
| FcZEB2-3N | TGGGCCGGGATACAAGTCT | |
| FcZEB2-3R | CAATGTTCTCTCATGGGAGGTAGTC | |
| FcZEB2 with 5F | GGAAAGTTCACGGTATAGCAAATC | |
| FcNRPS15-5F | AGAACCAATACCGAGACATAATGCT | *FcNRPS15* deletion mutant |
| FcNRPS15-5N | GTCAAATAGAGGCCCAACAAATG | |
| FcNRPS15-5R | GCACAGGTACACTTGTTTAGAGTCAGATTTTACTGGATTT AGGCTCG | |
| FcNRPS15-3F | CCTTCAATATCATCTTCTGTCGATTGTCAAATTACTCTGG TGAGGTT | |
| FcNRPS15-3N | TGCATTCTCCGGGGTCTTTT | |
| FcNRPS15-3R | ACGATGCTTTGGGGCGA | |
| FcNRPS15 with 5F | AAGCGGCGCTTATGACAGTG | |
| Gen with 5F | GTTGCCTAGTGAATGCTCCGTAACA | Geneticin screening |
| FgNRPS15com_F | TATAGGGCGAATTGGGTACTCAAATTGGTTAGCTTGATA GGAGAAAATACCAGAA | Complementation & Overexpression strain |
| FgNRPS15com_R | CCCGGTGAACAGCTCCTCGCCCTTGCTCACTCTCACTTTA TCTTCTCTCCGA | |
| FgNRPS15oe_F | TTTCGTAGGAACCCAATCTTCAAAATGGATACATGTCAA GATGATATTCTGT | |
| FgNRPS15oe_R | =FgNRPS15com_R | |
| NRPS15 seqF-1 | CAATAAGTGAGTTACTGCGCACAAAGC | *NRPS15* sequencing |
| NRPS15 seqR-1 | TTTACATGATCTGTGAAACGCTG | |

| | | |
|---|---|---|
| NRPS15 seqF-2 | CAGCAGTCTCAAGCGTGTAAGGAT | |
| NRPS15 seqR-2 | TACGCATTTCGCTCCCACAT | |
| NRPS15 seqF-3 | CCGAGCAAACAGAACAGTGACATA | |
| NRPS15 seqR-3 | TGGATCCAGAATACGCAACAG | |
| NRPS15 seqF-4 | CAAGGACGTTCACTGGGATTC | |
| NRPS15 seqR-4 | AGTCCCATGTCCCGTAATAGTCGT | |
| NRPS15 seqF-5 | CGTGGTTGTGGAGGTCGTCT | |
| NRPS15 seqR-5 | ATACGCCTGGTGCAATCTGTCC | |
| NRPS15 seqF-6 | CTACGTACACGAATTGTGGACAC | |
| NRPS15 seqR-6 | GTCTGATAGTGTTGTCCAGTTGCT | |
| NRPS15 seqF-7 | GAGTCCGATGAGCAAGGGGTTA | |
| NRPS15 seqR-7 | CTTTTGCAGCGGAGGGAGTTAT | |
| NRPS15 seqF-8 | TGGAACCATCAGGCACTATCTA | |
| NRPS15 seqR-8 | AGAGTCAATCTCTCTGCTGCTGC | |
| NRPS15 seqF-9 | AACGACGACCCCATACTTTTAGC | |
| NRPS15 seqR-9 | GGGTTGGTGGCCGCTATCT | |
| NRPS15 seqF-10 | TGGCATGAAAGTTATCTGTCTCGT | |
| NRPS15 seqR-10 | ACCTCCAGCAGATACTTGACATTT | |
| NRPS15 seqF-11 | TGGAGAACAAGGTGAATGACAA | |
| NRPS15 seqR-11 | AGTTCACCTTGATGCCGTTCTTC | |
| Rp27 with F | GGGAACTTTTTCTCACAGACACAA | |
| Native seqF-1 | CCAGCTGGCGAAAGGGG | |
| NRPS15 pyes2_F(hind3) | aaaagcttATGGATACATGTCAAGATGATATTCTGTC | *NRPS15* amplification for pYES2 cloning |
| NRPS15 pyes2_R(Xho1) | aaactcgagTTATCTCACTTTATCTTCTCTCCGAA | |
| pYES2 seqF | TAACGTCAAGGAGAAAAAACCCC | pYES2 sequencing |
| pYES2 seqR | CCTTCTCAAGCAAGGTTTTCAGTA | |
| zeb1 qPCR-F | CGTTCATATCGTCTCTCACATCCA | RT-qPCR |
| zeb1 qPCR-R | AGTTGGGTACAACACTGCGGG | |
| pks4 qPCR-F | CATCCAAACAGCAAACGACAAAA | |
| pks4 qPCR-R | GCAACCTGTTTGAGGGGCTTA | |
| pks13 qPCR-F | TCTCTTCGGAGATCAAACTGACTC | |
| pks13 qPCR-R | CTGGAGATTTGCAAAGTCGCC | |
| zeb2 qPCR-F | ATGCAATCTACCGAGTCGTTCCAT | |
| zeb2 qPCR-R | ATTCGACGTTTGGCATTGGTTGTTGAC | |
| zeb2L qPCR-F | =zeb2 qPCR_F | |
| zeb2L qPCR-R | =zeb2 qPCR_R | |
| zeb2S qPCR-F | AGTCGTATTCTCAAGGTTATTCAGGATG | |
| zeb2S qPCR-R | ATATCTCTTGTTGGTAATCATGTGCTTCA | |
| cyp1 qPCR-F | TCAAGCTCAAGCACACCAAGAAGG | |

| | |
|---|---|
| cyp1 qPCR-R | GGTCCGCCGCTCCAGTCT |
| BY4741 act_F | CTATTGTTGACCGTCGAAGA |
| BY4741 act_R | AAGTGCCAAGATAGAACCAC |

98
